# Quinolinate Promotes Immune Tolerance through Macrophage Polarization in Glioblastoma

**DOI:** 10.1101/2022.07.12.499609

**Authors:** Pravin Kesarwani, Shiva Kant, Yi Zhao, Prakash Chinnaiyan

## Abstract

There has been considerable scientific effort dedicated to understanding the biologic consequence and therapeutic implications of aberrant tryptophan metabolism in brain tumors and neurodegenerative diseases. An overwhelming majority of this work has focused on the upstream metabolism of tryptophan; however, this has not resulted in clinical application. Using global metabolomic profiling of patient-derived brain tumors, we identify the downstream metabolism of tryptophan and accumulation of quinolinate (QA) as a metabolic node in glioblastoma and went on to demonstrate its critical role in promoting immune tolerance. QA acts as a “metabolic checkpoint” in glioblastoma by inducing NMDA receptor activation and Foxo1/PPARγ signaling, resulting in amplification of immune suppressive macrophages. Using a genetically-engineered mouse model designed to inhibit production of QA, we identify kynureninase as a promising therapeutic target to revert the potent immune suppressive microenvironment in glioblastoma. These findings offer the scientific community an opportunity to revisit the biologic consequence of this pathway as it relates to oncogenesis and neurodegenerative disease and a framework for developing new immune modulatory agents to further clinical gains in these otherwise incurable diseases.

## INTRODUCTION

The capacity of cancer cells to evolve mechanisms to evade the host immune system represents a requisite event for these cells to survive and continue to proliferate and metastasize.^1^ Accordingly, investigations have identified numerous ways by which these cells actively sculpt the tumor microenvironment to promote immune tolerance. In brain tumors, tumor-associated macrophages (TAMs) have emerged as critical cells contributing towards their potent immune suppressive microenvironment.^2^ Macrophages and microglial cells (resident macrophages) constitute over 30% of immune fraction in gliomas and, therefore, act as major component of neuroinflammation and immune dysregulation in glioblastoma (GBM).^3, 4^ Earlier research has classified TAMs as proinflammatory macrophages (M1 macrophages), responsible for anti-tumorigenic function and anti-inflammatory macrophages (alternatively activated M2 macrophages) responsible for pro-tumorigenic capacity. Alternatively activated M2 macrophages are a type of TAM contributing towards immunosuppression, angiogenesis, and tumor progression and, therefore, an important target for immunotherapy.^5–7^

Recent investigations have uncovered elegant strategies that cancer cells use to promote a state of immune tolerance in the tumor microenvironment by exploiting conserved, metabolic signaling.^8–11^ Aberrant tryptophan metabolism, and its metabolic intermediate kynurenine (Kyn), represents one such pathway that has received considerable attention (**Fig. 1A**). Tryptophan is metabolized to Kyn by the rate limiting enzymes indoleamine 2,3-dioxygenase 1 (IDO1) and tryptophan 2,3-dioxygenase (TDO)^12, 13^ and contributes to an immune tolerant environment at many levels. Intriguingly, the most notable physiologic role of this metabolic pathway has been attributed to peripheral immune tolerance and fetal protection from maternal immune rejection in the placenta.^14^ A variety of tumors, including GBM, have evolved mechanisms to co-opt this potent mechanism of suppression to evade the host immune system, which has been an active area of research with a particular focus on the immune-modulatory metabolite Kyn.^12, 13, 15^

**Figure 1.**
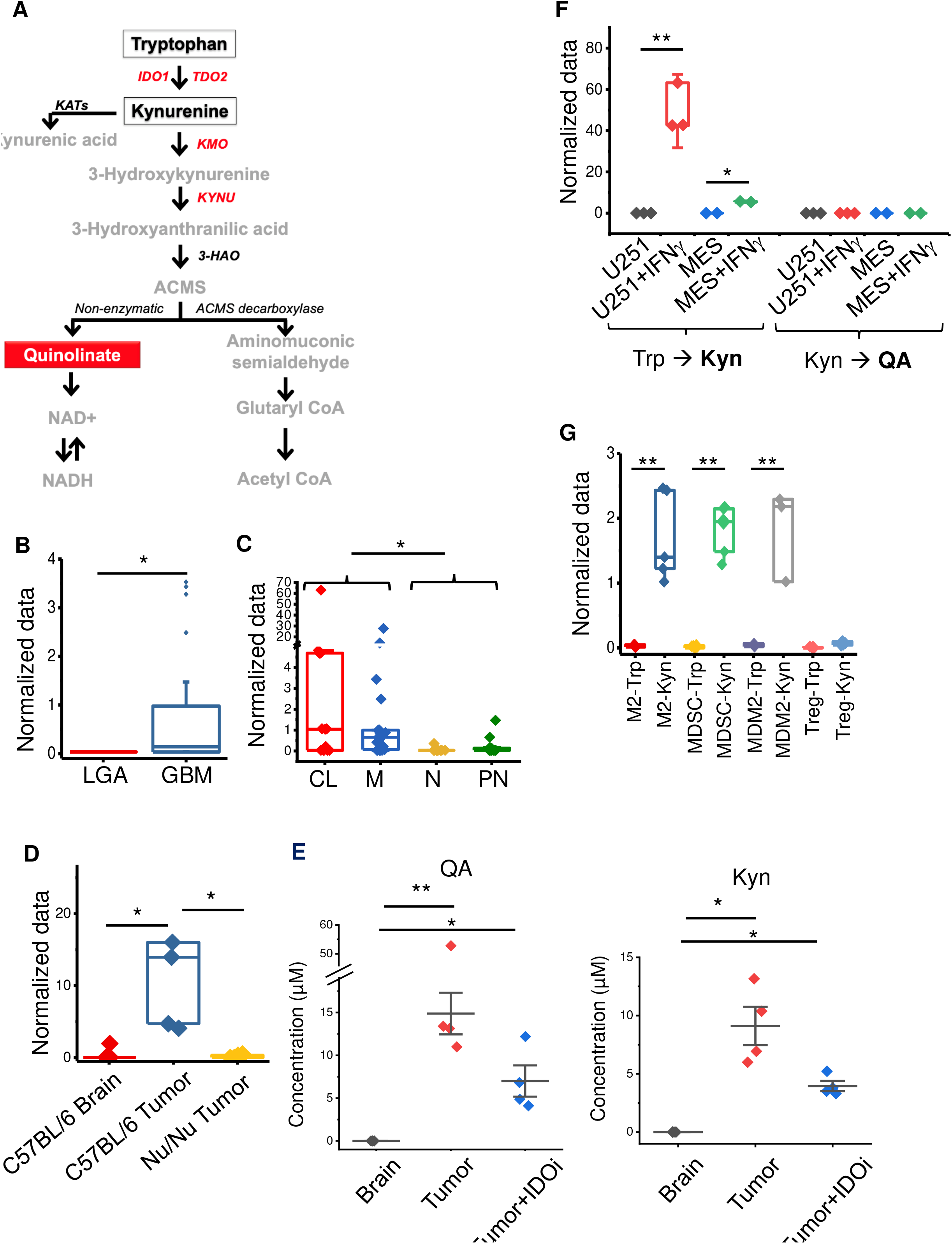
Quinolinate (QA) accumulation in glioblastoma (GBM). (**A**) Schematic of tryptophan (Trp) metabolism. (**B**) Metabolomic profiling performed on patient-derived low-grade astrocytoma (LGA; n=28) and GBM (n=80) demonstrates differential accumulation of QA in GBM. (**C**) The relative accumulation of QA was evaluated in molecular subtypes of GBM (n=56), classified as classical (CL), mesenchymal (M), neural (N), or proneural (PN). (**D**) The murine GBM line TRP was grown orthotopically in C57BL/6 mice and analyzed for QA and compared to normal murine brain, and human-derived GBM cells (MES line) grown in Nu/Nu mice (n=6/group). (**E**) QA and kynurenine (Kyn) were quantified in normal brain and TRP tumors grown orthotopically in C57BL/6 mice +/- the IDO inhibitor (IDOi) GDC-0919 (n=4/group). (**F**) The upstream and downstream metabolism of tryptophan (Trp→Kyn and Kyn→QA, respectively) was evaluated in the described cell lines +/- IFNγ (n=2-3/group). (**G**) M2 macrophages, myeloid-derived-suppressor cells (MDSC), microglial-derived M2 cells (MDM2), and regulatory T cells (Treg) were isolated from C57BL/6 mice, cultured with Trp or Kyn, and analyzed for QA (n=5/group). Box plots represent interquartile range, line between data points represents mean, and whiskers represent SE. *=p<0.05; **=p<0.005

Although, accumulation of quinolinate (QA) has been both recognized and a focus of neuropathology for nearly 4 decades, its role in shaping the immune microenvironment remains unknown.^16–18^ Specifically, as QA does not cross the blood brain barrier, concentrations are typically low (<100 nM) in the brain, however, its accumulation has been observed in a variety of pathologic states, including Alzheimer’s and Huntington’s disease, acquired immunodeficiency syndrome (AIDS) dementia complex, trauma, and meningitis.^19–22^ Early studies in neuropathology demonstrated the accumulation of QA leads to neuronal death/dysfunction through several mechanisms, including N-methyl-D-aspartate (NMDA) receptor-mediated excitotoxicity, lipid peroxidation, and synthesis of nitric oxide.^23–25^. However, its capacity to modulate other biological functions remains largely undefined. Recently, our group has identified the accumulation of QA in GBM, suggesting a contributory role in tumor growth.

In this study, we discovered the accumulation of QA to be an important “metabolic checkpoint” regulating immunity in GBM. We uncovered a novel mechanism by which QA utilizes the NMDA-receptor to induce Foxo1/PPARγ transcription factor signaling, exacerbating M2 macrophages into a highly immune-suppressive state. Targeting this pathway demonstrated anti-tumor activity, offering strong promise in GBM and a framework that may be applied for the treatment of a variety of neurodegenerative diseases where QA has been demonstrated to accumulate.^19, 26, 27^

## RESULTS

### Quinolinate accumulation in glioblastoma

In our previous work, we performed cross-platform analyses coupling global metabolomic profiling with genomics in over 100 patient-derived brain tumors to provide a window into specific metabolic programs driving the aggressive phenotype of this malignancy in the context of its molecular architecture.^9, 28^ Through these studies, we identified the accumulation of QA, the downstream metabolic intermediate of tryptophan, in GBM, with a ~64-fold increase when compared to low-grade astrocytoma (LGA; **Fig. 1B**). We went on to determine if QA accumulation was a direct consequence of established molecular subtypes in GBM. Although outliers with particularly high levels of QA were only observed in isocitrate dehydrogenase 1 (IDH1) wild-type (WT) and O-6-methylguanine-DNA methyltransferase (MGMT) unmethylated tumors, two of the strongest prognostic factors in GBM,^29, 30^ collectively, levels of this metabolite were not significantly different between these subtypes (**Fig. S1**). Next, using gene expression profiles generated from these matched samples, tumors were classified according to their molecular subtype: mesenchymal, proneural, classical, or neural.^31^ Interestingly, the accumulation of QA was specific to the mesenchymal and classical subtypes, when compared to proneural and neural subtypes (**Fig. 1C**). The accumulation of QA in GBM was validated in an independent data set using immunohistochemical (IHC) staining (**Fig. S2**).

We next sought to determine if QA accumulation was recapitulated in GBM preclinical models. QA was present when evaluated in cell lines derived from a genetically engineered mouse model (GEMM ^32, 33^ grown intracranially in C57BL/6J mice, but not in normal brain (**Fig. 1D)**, which we went on to quantify (mean concentration ~20 μM; **Fig. 1E**). Interestingly, QA was not detected when evaluated in patient-derived tumors grown in immune deficient mice (**Fig. 1D)**, suggesting immune cells may play a role in the intermediary metabolism of tryptophan and the accumulation of QA. As discussed above, targeting the upstream metabolism of tryptophan has been an active are of investigation. We therefore, sought to determine how targeting the upstream metabolism of tryptophan may influence the accumulation of the downstream metabolic intermediate QA.

Using the IDO inhibitor GDC-0919, we demonstrate that although a reduction is Kyn is observed, it is not complete (**Fig. 1E**), which is likely attributed to parallel pathways a tumor may utilize to generate this metabolite independent of IDO1. Importantly, in a similar fashion, when evaluating for QA, although IDO inhibition resulted in a decrease in QA, biologically relevant concentrations were maintained in the tumor, suggesting a more specific, downstream enzyme may be a more relevant target to attenuate the accumulation of QA.

Although we and others have previously demonstrated that upon IFN-γ mediated activation, GBM cell lines are able to generate Kyn when cultured with tryptophan,^10, 13^ here we show that these cells do not have the capacity to independently complete this metabolic pathway in its entirety, as QA was not detected when cultured with either tryptophan or Kyn (**Fig. 1F**). As our above investigations suggested immune cells may play a role in the accumulation of QA, we evaluated the potential of immune cells typical to the GBM microenvironment, including M2 macrophages, myeloid derived suppressive cells (MDSCs), microglia derived M2 macrophages (MDM2s), and regulatory T cells (Tregs), to generate QA. Interestingly, paradoxical to tumors, although myeloid derived immune cells and microglia did not have the capacity to metabolize tryptophan, they were able to metabolize Kyn to generate QA (**Fig. 1G**). Collectively, the accumulation of QA represents a metabolic phenotype of GBM and is a direct consequence of a dynamic interaction between tumor and immune cells.

### QA contributes towards immune suppression by priming macrophage polarization towards the M2 phenotype

Next, we sought to determine the biologic consequence of QA in GBM. As the role of QA’s upstream mediator Kyn in immune response has been well-described,^10, 12–14^ we evaluated for possible immune consequences of QA. As an initial investigation, we examined the potential for QA, along with its upstream and downstream metabolic intermediates, Kyn and NAD+ respectively (**Fig. 1A**), to polarize and/or promote proliferation of immune suppressive cells. Of the immune cells tested, Tregs, MDSCs, and macrophages, only macrophages appeared to be influenced by QA, resulting in a dramatic increase in polarization from 47.4±5.3% to 70.9±7.8% (**Fig. 2A**). This was not observed with Kyn or NAD+, suggesting direct action of this metabolite rather than the consequence of its downstream metabolism. We went on to extend these novel findings by demonstrating the ability of QA to increase expression of the M2 macrophage specific markers CD206 and IL4Rα, a suppressive marker of M2 macrophages ^34, 35^ (**Fig. 2B**). As we have already established that macrophages have the unique ability to metabolize Kyn to QA, we hypothesized that rather than the standard 7 d schedule of polarization, a longer schedule (9-10 d) would provide these cells time to generate QA, which could in turn, could influence macrophage plasticity. As predicted, although not as potent as QA, a longer schedule with Kyn did result in increased macrophage polarization. Importantly, these changes were mitigated by inhibiting the upstream biosynthetic enzyme kynurenine-3-monooxygenase (KMO; UPF648), in Kyn, but not in QA treated cells (**Fig. 2C**), further supporting the direct role QA plays in macrophage polarization. The potential for QA to modulate macrophage polarization was validated in the established murine macrophage cell line IC-21 (**Fig. S3**).

**Figure 2.**
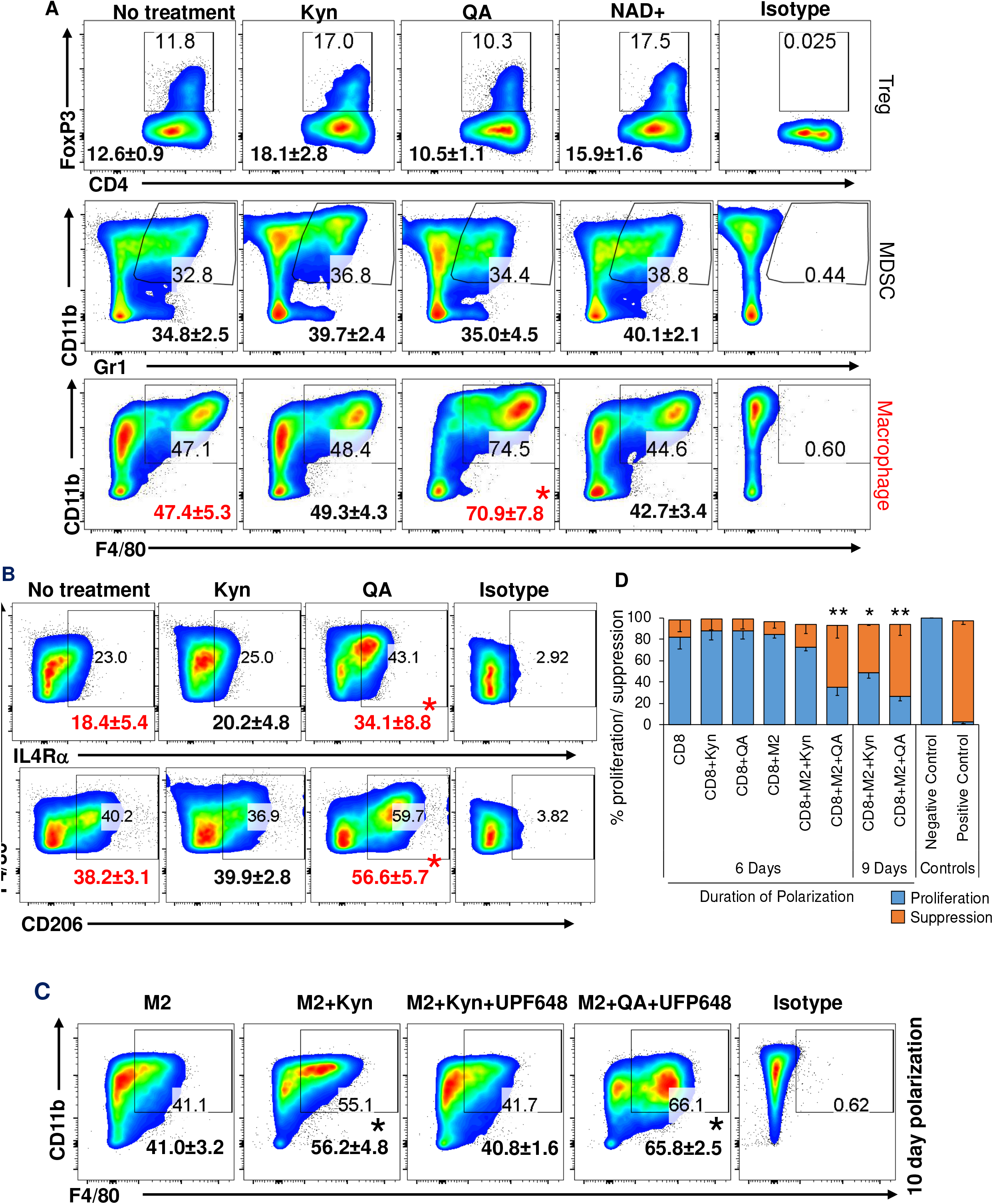
QA contributes towards immune suppression by priming monocyte polarization towards the M2 phenotype. (**A**) Regulatory T cells (Treg; CD4+FoxP3+), myeloid derived suppressor cells (MDSC; Gr1+CD11b), and macrophages (CD45+F4/80+CD11b+) were obtained from C57BL/6 mice and cultured in ± kynurenine (Kyn-20μM), QA (20 μM), or NAD+ (10μM) and analyzed by flow cytometry. (**B**) Macrophages polarized towards the M2 phenotype ± Kyn or QA (20 μM) were gated for CD45+F4/80+CD11b+ and analyzed for CD206 or IL-4Rα using flow-cytometry. (**C**) Similar studies were performed on macrophages polarized towards the M2 phenotype ± Kyn or QA for an additional 3 d (10 d total, to allow for further metabolism/generation of QA) ± the kynurenine 3-monooxygenase (KMO) inhibitor UFP648. (**D**) To analyze the functional suppression of M2 macrophages, M2 cells were polarized ± Kyn or QA (20 μM) for 6 or 9 d. Splenocytes from C57BL/6 mice were used for isolating CD8+ T cells using magnetic bead sorting. CFSE labeled CD8+ T cells were activated using plate-bound anti-CD3/CD28 antibody for three days in the presence or absence of M2 cells ± Kyn or QA. Data is represented as a bar graph demonstrating CFSE dilution (proliferation) or suppression. CFSE labeled CD8+ T cells without stimulation (anti-CD3/CD28 antibody) were used as a positive control of suppression. Unlabeled CD8+ T cells were used as a negative control for proliferation. Data is representative of a minimum of three experiments. Numbers represent mean±SD. Bar graph shows mean±SD. *=p<0.05; **=p<0.005

We went on to determine if the observed changes in M2 macrophage polarization translated to the capacity of these cells to functionally suppress CD8+ T-cell proliferation using CD8/M2 macrophage co-culture experiments. At the indicated CD8:M2 ratio (5:1), M2 macrophages and QA alone did not inhibit CD8 cell proliferation (**Fig. 2D, S4A**). Consistent with polarization studies, M2 macrophages cultured in the presence of QA led to potent inhibition of CD8+ proliferation (35.1±7.6%), while Kyn resulted in minimal changes (72.4±2.9%) when tested in standard 6 d polarization conditions. When polarization was extended to 9 d (allowing time for M2 macrophages to metabolize Kyn to QA), inhibition of CD8+ proliferation was observed (48.3±5.2%), albeit less than when cultured with QA (26.7±4.2%). In addition to CD8+ proliferation, QA significantly their expression of granzyme B+ CD8 cells (**Fig. S4B**) and its influence on proliferation were validated when extended to mouse derived microglia cells (**Fig. S5A/B**).

### QA confers an “M2-like” phenotype to M1 macrophages and microglia

As we identified a novel role QA plays in polarizing macrophages towards an M2 phenotype with immune suppressive properties, we extended investigations to determine if this metabolite influenced M1 macrophage polarization and/or phenotype. Although QA did not appear to influence M1 polarization (CD45+CD11b+F4/80+CD80hi, **Fig. S6A**), intriguingly, it did confer an “M2-like” molecular phenotype to these cells, with the percentage of M1 macrophages expressing the M2 macrophage marker CD206+ dramatically increasing from 5.4±2.9% to 25.3±8.6% (**Fig. 3A**). Even more striking, in addition to conferring an “M2-like” molecular phenotype, M1 macrophages cultured in the presence of QA gained the ability to suppress T-cell proliferation (from 76.8±3.2% to 54.1±2.1%; **Fig. 3B**), thereby, functionally mimicking M2 macrophages. As a key function of M1 macrophages is phagocytosis, we went on to determine if QA-induced skewing of macrophage polarization could result in a functional consequence on phagocytosis efficiency. As expected, murine derived microglia polarized towards the M1 phenotype demonstrated the unique ability of phagocytosis when compared to cells polarized towards the M2 phenotype, as measured by fluorescently labeled β-amyloid peptide (**Fig. 3C**; 92.2% vs. 18.7%). Intriguingly, consistent with molecular studies, QA conferred an “M2-like” phenotype to M1 macrophages, significantly attenuating their phagocytosis efficiency (38.8%). Importantly, these findings in mouse lines were further validated in human microglia derived from induced pluripotent stem cells (iPSCs; iCell; **Fig. 3D**) and murine-derived M1 macrophages (**Fig. S6B**).

**Figure 3.**
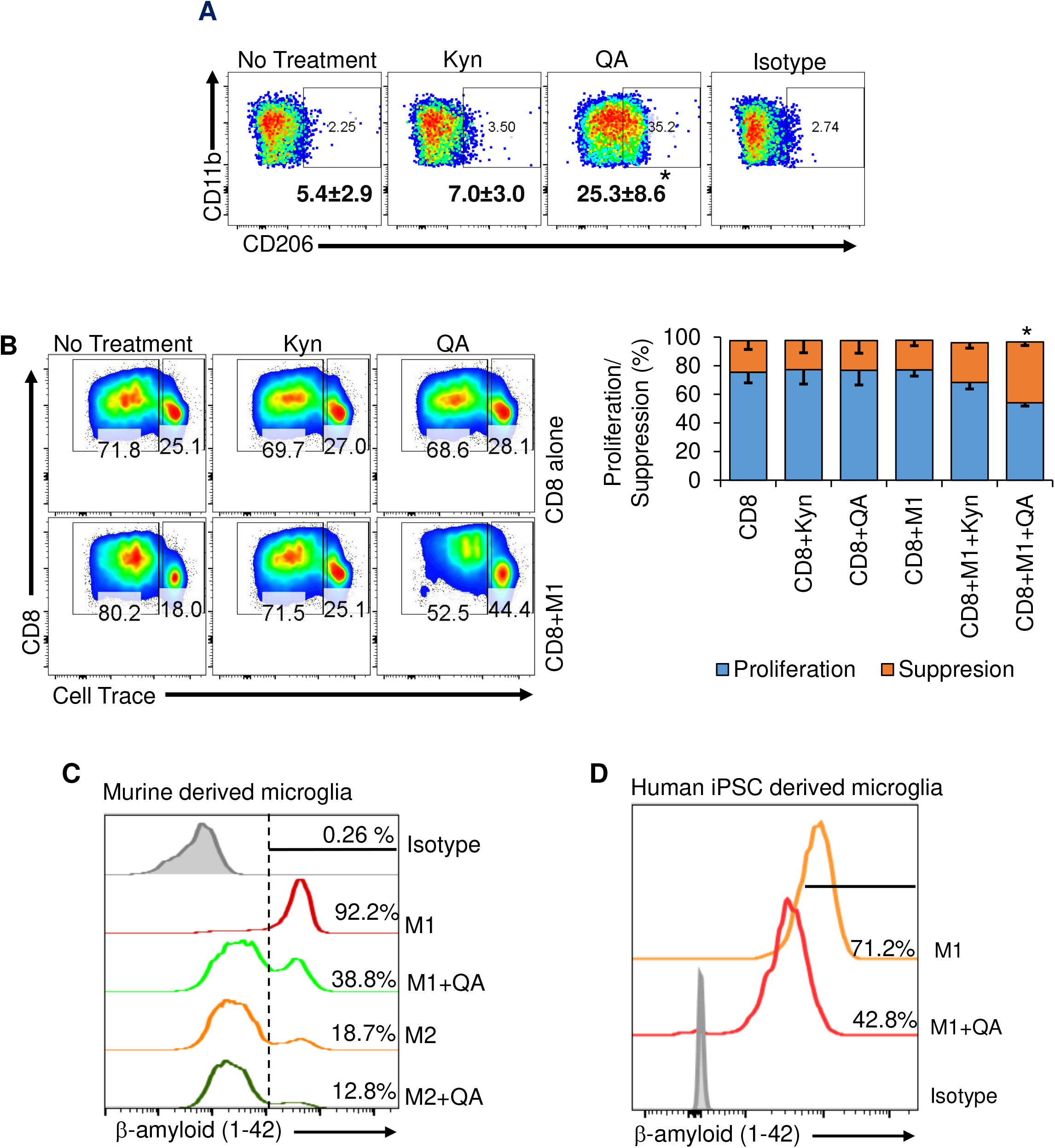
QA induces an “M2-like” phenotype in M1 macrophages and microglia. (**A**) M0 macrophages harvested from the bone marrow of C57BL/6 mice were polarized to M1 macrophages with LPS (100ng/ml) and IFNγ (50ng/ml) for 24-36 h ± Kyn or QA (20 μM) and gated for M2 macrophage markers (CD45+F4/80+CD11b+CD206+). Numbers represent mean±SD. (**B**) The functional suppression of M1 macrophages polarized +/- Kyn or QA was performed using CD8/macrophage co-culture experiments with CFSE dilution, as described in Fig. 2. (**C**) Murine-derived microglia and (**D**) human induced pluripotent stem cell (iPSC) derived microglia were matured in the presence of GM-CSF conditioning media and polarized towards the M1 (LPS+IFNγ) or M2 (IL4+IL13) phenotype ± QA (20 μM). Cells were pulsed with green florescent β-amyloid (1-42) peptide and analyzed for phagocytosis of this peptide at 16 h by flow cytometry. Data is representative of a minimum of two experiments. Bar graph shows mean±SD. *=p<0.05.

### QA modulates M2 macrophage polarization through the NMDAR/Foxo1/PPARγ signaling axis

As QA’s upstream metabolic intermediary Kyn has been shown to modulate immune response through aryl hydrocarbon receptor (AhR) activation,^12^ as an initial investigation, we sought to determine if this transcription factor served as a potential target and underlying mechanism of QA-induced modulation of macrophage plasticity. Utilizing cells transfected with a luciferase reporter functionally linked to the AhR-response promoter, we confirmed that Kyn, but not QA, has the unique ability to activate the AhR (**Fig. S7A**). To provide a framework in identifying mechanistic underpinnings linking QA with M2 macrophage polarization, we defined global changes of QA-induced gene expression in M2 macrophages. In these studies, M2 macrophages (CD45+CD11b+F4/80+CD206+) cultured ± QA were flow-sorted, and mRNA was isolated and profiled using an Affymetrix mouse array (**Fig. 4A**). To begin to understand the signaling machinery orchestrating QA-induced M2 macrophage polarization, Pathway Commons was used to map protein-to-protein interactions between the top 25 differentially expressed genes identified by volcano plot (**Fig. 4B**), identifying peroxisome proliferator-activated receptor gamma (PPARγ), which has been previously linked to M2 macrophage polarization,^36, 37^ as a lead candidate (**Fig. 4C**).

**Figure 4.**
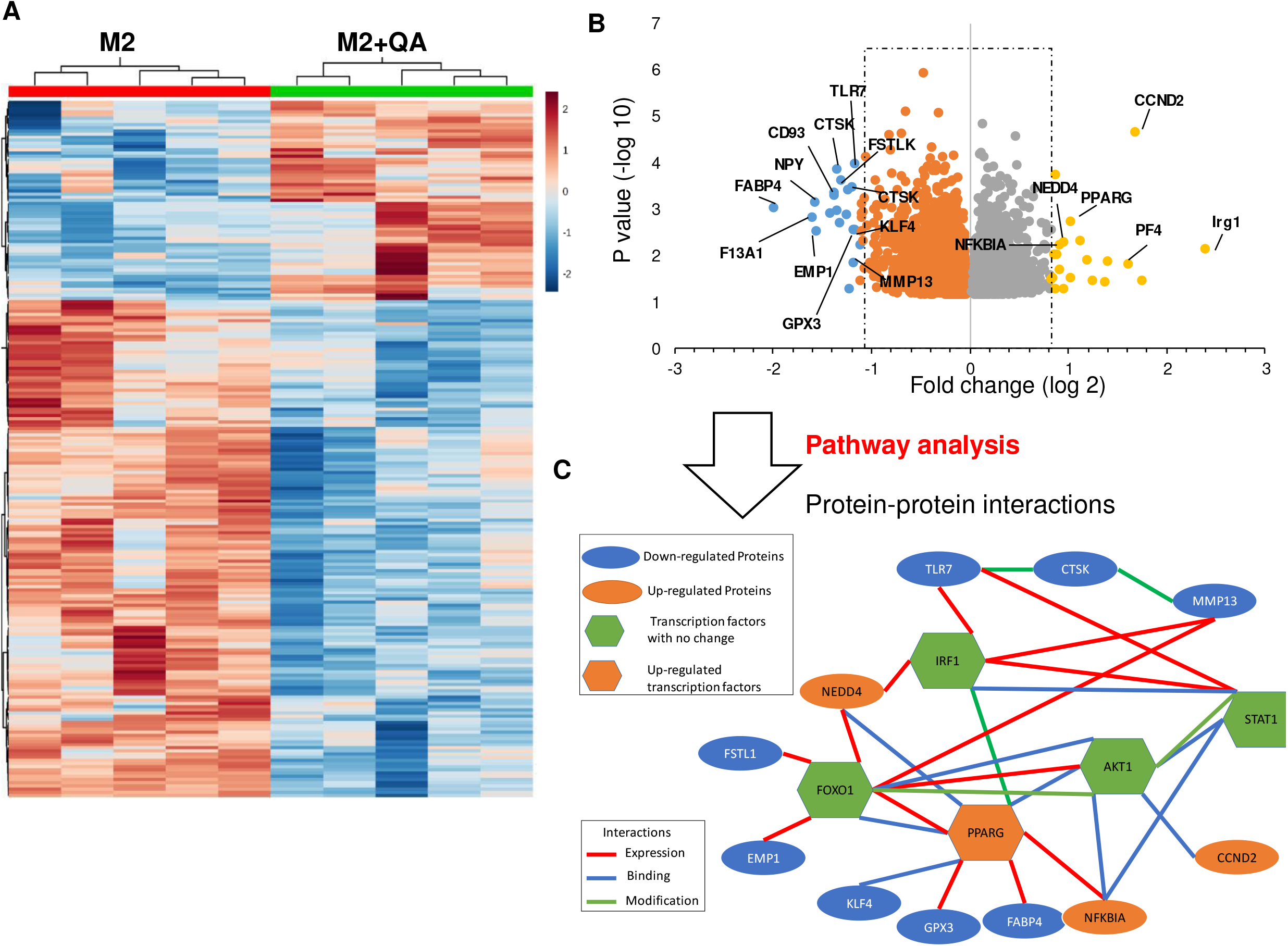
QA modulates PPARγ mediated transcriptional programs in M2 macrophages. M2 macrophages cultured in presence or absence of QA (20 μM) were flow-sorted for M2 macrophages (CD45+CD11b+F4/80+CD206+), and mRNA analyzed using Affymetrix arrays (Clariom™ D Assay, mouse; n=5 samples /group). (**A**) Heatmap demonstrates differentially expressed genes (log 1.5-fold change). (**B**) Differentially expressed genes (p<0.05) are presented as a volcano plot. (**C**) Protein-protein interaction and pathway analysis.

To extend transcriptomic findings, we evaluated for differential expression of PPARγ in our models. Consistent with these studies, M2 macrophages cultured in the presence of QA demonstrated increased expression of PPARγ (**Fig. 5A**). As expected, culturing M2 macrophages in the presence of the PPARγ agonist troglitazone demonstrated a similar increase in PPARγ expression, and importantly, the combination of QA and troglitazone did not lead to any further increase in expression, suggesting a commonality in underlying mechanisms (**Fig. S7B**). Next, we extended these findings to evaluate for M2 macrophage polarization. Consistent with western blot data, troglitazone increased M2 macrophage polarization in a similar manner as QA, however, no further increases were observed with the combination, again supporting a commonality in underlying mechanisms (**Fig. 5B**). To further validate this pathway, M2 macrophages were polarized in the presence of the PPARγ antagonist GW9662. As expected, this led to decreased levels of M2 macrophages and was no longer influenced by QA, further supporting the hypothesis that QA modulates M2 macrophage polarization through PPARγ, which we went on to molecularly validate in the mouse macrophage IC-21 line using siRNA against PPARγ (**Fig. S7C**). Lastly, to directly determine if QA activates PPARγ-dependent transcriptional programs in M2 macrophages, we utilized a PPARγ specific transcriptional activation assay, further supporting the role QA plays in modulating PPARγ transcriptional activation (**Fig. 5C**).

**Figure 5.**
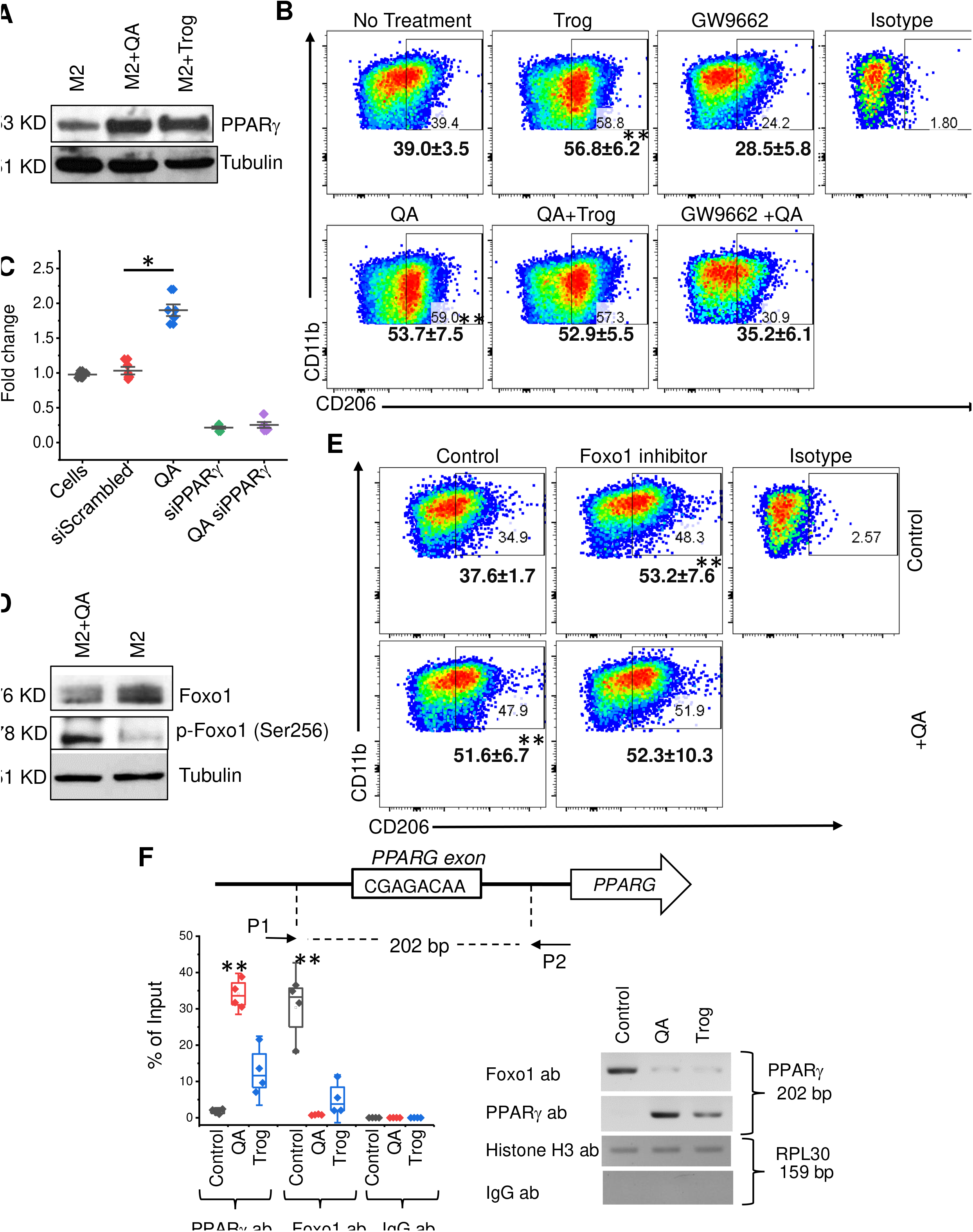
QA regulates PPARγ signaling using Foxo1. (**A**) Macrophages obtained from C57BL/6 mice were polarized towards the M2 phenotype ± QA (20 μM) or the PPARγ agonist troglitazone (Trog; 5 μM) and evaluated for the indicated proteins by western blot. (**B**) Macrophages obtained from C57BL/6 mice were polarized towards the M2 phenotype ± Trog, GW9662 (PPARγ antagonist) and/or QA (20 μM) and analyzed for M2 macrophages (CD45+CD11b+F4/80+CD206+) by flow cytometry. Numbers represent mean±SD. (**C**) PPARγ transcriptional activity was evaluated in the murine macrophage cell line IC-21 polarized towards the M2 phenotype ± siRNA of indicated proteins by PPARγ transcriptional activity ELISA (n=4). (**D**) C57BL/6 mouse-derived macrophages were polarized towards the M2 phenotype ± QA (20 μM) and evaluated for the indicated proteins by western blot. (**E**) Macrophages were cultured in the presence or absence of QA ± the Foxo1 inhibitor AS-1842856 (0.1μM) and evaluated for M2 macrophages (CD45+CD11b+F4/80+CD206+) by flow cytometry. Numbers represent mean±SD. (**F**) Chromatin immunoprecipitation (ChIP) was performed on C57BL/6 mouse-derived macrophages polarized towards the M2 phenotype ± Trog or QA. ChIP was performed using antibodies against PPARγ, Foxo1, and histone 3 (positive control), and IgG antibody (negative control). After IP and DNA purification, the PPARγ exon (202 bp in length) was analyzed using qPCR and gel electrophoresis. Box plots represent interquartile range, line between the data points represents mean, and whiskers represents SE. *=p<0.05; **=p<0.005

We next sought to determine how QA leads to PPARγ activation. Re-analyzing transcriptional profiles generated from QA-induced M2 macrophages and specifically evaluating for known regulators of PPARγ, the transcriptional factor forkhead box protein O1 (Foxo1) emerged as a putative molecular mediator. The Foxo1 transcription factor plays key roles in several cellular processes, including proliferation, metabolism, and survival in various immune cells. Several studies have demonstrated that Foxo1 inhibits the activity of PPARγ by binding to its promoter and inhibiting transcription,^38, 39^ or by directly binding to PPARγ protein in a transrepressional manner, resulting in the inability of PPARγ to bind to its primary locus, the PPARγ response element (PPRE).^40^ Transrepressional activity is blunted by phosphorylation of Foxo1, inducing cytosolic localization and ubiquitination, leading to PPARγ activation.^39^ We therefore explored the interface between QA and Foxo1 in further detail. Consistent with our working model, M2 macrophages cultured in the presence of QA led to diminished expression of the negative regulator of PPARγ, Foxo1, and its phosphorylation at Ser256, signaling ubiquitination of this protein (**Fig. 5D)**. These molecular findings were further validated using the Foxo1 inhibitor AS1842856, which increased M2 macrophage polarization in a manner similar to PPARγ activation and was no longer influenced by QA (**Fig. 5E)**. Lastly, chromatin immunoprecipitation (ChIP) was performed, definitively establishing that QA attenuates Foxo1 binding to the PPARγ promotor in a manner similar to the PPARγ agonist troglitazone (**Fig. 5F**). Interestingly, these studies uncovered a strong QA-induced positive feedback loop involving PPARγ expression and activation of its own promoter.

As we have established that QA modulates M2 macrophage polarization through the Foxo1/PPARγ axis, we next sought to determine the primary target of QA. As QA is a known ligand of the NMDA receptor, as an initial investigation, we evaluated for the expression of the NMDA receptor in macrophages. Interestingly, NMDAR1 expression (a key receptor subunit of this heterotetramer) was only observed in M2 macrophages but not unpolarized macrophages (**Fig. 6A**). Similar to QA, an increase in M2 macrophage polarization was observed when cultured in the presence of NMDA and consistent with our working model, the NMDAR competitive inhibitor L-AP5 demonstrated a dose-dependent inhibition of M2 macrophage polarization and mitigated the activity of QA (**Fig. 6B**). Lastly, we demonstrated that NMDA led to Foxo1 phosphorylation and increased PPARγ expression in a manner similar to QA and troglitazone (**Fig. 6C**). Collectively, we demonstrate QA-induced M2 macrophage polarization is mediated through the NMDAR/Foxo1/PPARγ signaling axis (**Fig. 6D**).

**Figure 6.**
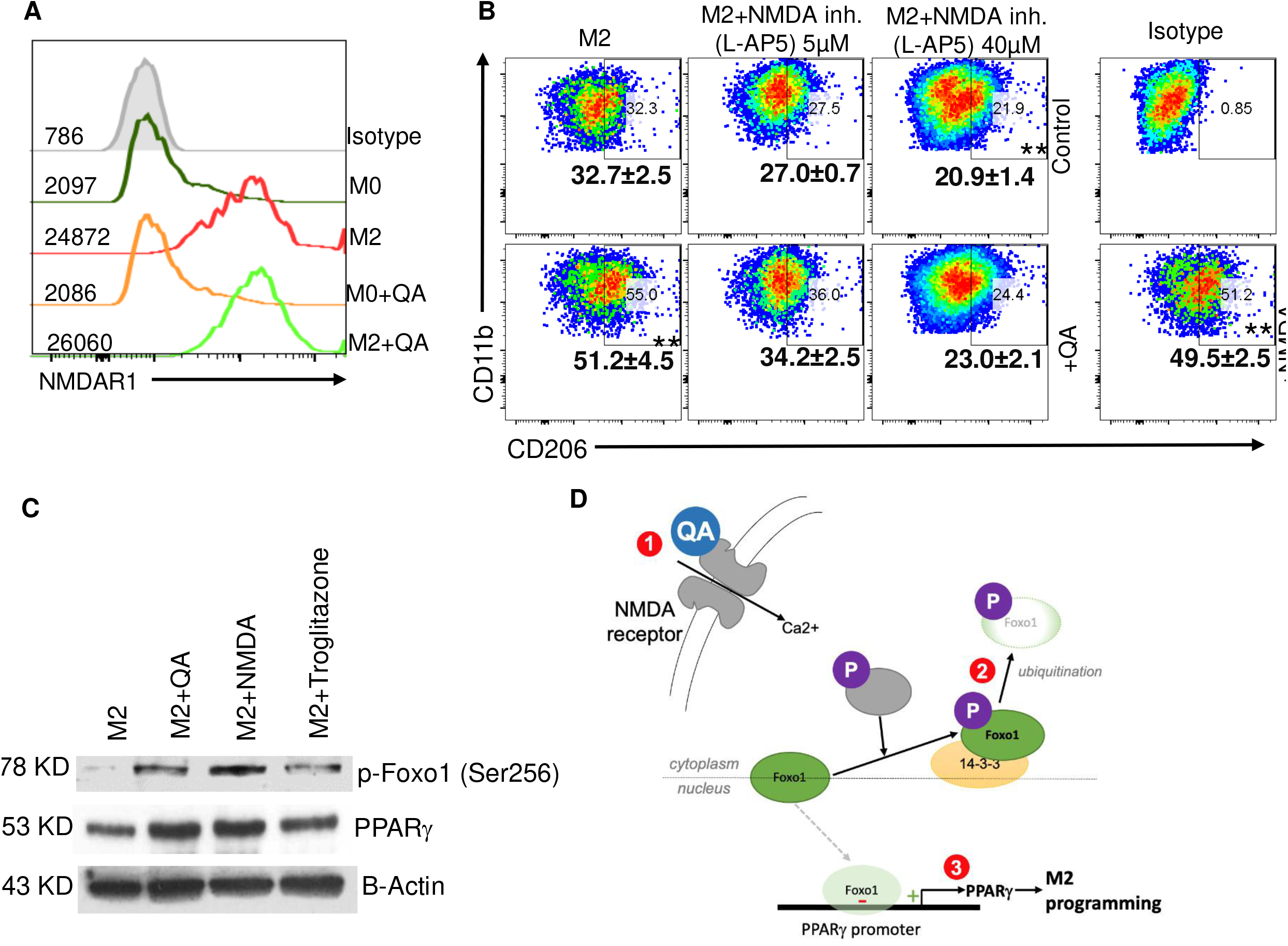
QA modulates M2 macrophage polarization through the NMDAR/Foxo1/PPARγ signaling axis. (**A**) M2 macrophages were polarized ± QA, gated on the macrophage marker (CD45+CD11b+F4/80+) and CD206+ve or CD206-ve, and evaluated for expression of the NMDA receptor 1 (NMDAR1). (**B**) M2 macrophages were cultured in ± QA, L-AP5 (NMDAR1 inhibitor), or NMDA. Cells were gated on the M2 macrophage marker (CD45+CD11b+F4/80+ CD206+). (**C**) M2 macrophages were cultured in ± QA, NMDA, or Trog, and indicated proteins were evaluated by western blot. (**D**) Schematic depicting the proposed mechanism of QA-induced M2 macrophage polarization. QA binds to the NMDA receptor of macrophages *(1),* leading to phosphorylation of Foxo1 *(2).* Phosphorylated Foxo1 is retained in the cytoplasm and destined for ubiquitination. Loss of nuclear Foxo1, a negative regulator of the PPARγ promoter, leads to increased PPARγ expression and transcriptional programs designed to promote macrophage polarization towards the M2 phenotype *(3).* Data is representative of a minimum of three experiments. Numbers represent mean±SD. *p<0.05; **=p<0.005.

### Targeting the downstream metabolism of tryptophan in GBM by inhibiting KYNU

The above findings support targeting the downstream metabolism of tryptophan may serve as a novel therapeutic strategy to overcome M2 macrophage-induced immune suppression in GBM. As the primary cell able to generate QA from tumor-produced Kyn are macrophages, we evaluated for the expression of enzymes involved in the downstream metabolism of tryptophan in mouse derived macrophages using RT-PCR. Baseline levels of KMO, kynureninase (KYNU), and 3-HAO (3-hydroxyanthranilate 3,4-dioxygenase) appeared equivalent. However, interestingly, when cultured in the presence of QA, there appeared to be a strong positive feedback loop in all enzymes, but most notable in KYNU, which demonstrated ~1000x increase in expression (**Fig. 7A**). Based on these findings, KYNU emerged as our primary therapeutic target to inhibit QA accumulation in GBM. As agents effectively targeting this enzyme are not commercially available, we generated a *Kynu*-/- GEMM to allow us to extend these promising findings *in vivo.* C57BL/6NJ-Kynuem1J/Mmjax mice were developed using CRISPR-Cas9 knockout of *Kynu* exon 3 and 219bp flanking intronic sequences and F3 (homozygous to homozygous mating). Successful knockout was validated using qPCR (**Fig. S8A)**. No clear phenotypic changes were observed in *Kynu-/-* mice (**Fig. 7B**). We further validated KYNU expression was abrogated in both M2 and M1 macrophages derived from *Kynu-/-* mice by western blot (**Fig. 7C**) and the inability of Kyn to modulate polarization of these cells (**Fig. S8B**). Next, we sought to validate cells contributing towards immune response remained intact in the *Kynu-/-* model. CD4+ T cells, CD8+ T cells, Tregs, B cells and NK cells generated from splenocytes of WT and *Kynu*-/- mice were quantitatively similar (**Fig. S8C**) and CD4+ and CD8+ T cells demonstrated similar phenotypes upon anti-CD3/anti-CD28 activation, including proliferation (Ki67), TNFα and IFNγ inflammatory cytokine production, and T cell activation marker CD69 (**Fig. S8D**). Further QA, but not Kyn, retained the ability to induce functional suppression in M2 macrophages derived from *Kynu*-/- mice (**Fig. S8E**).

**Figure 7.**
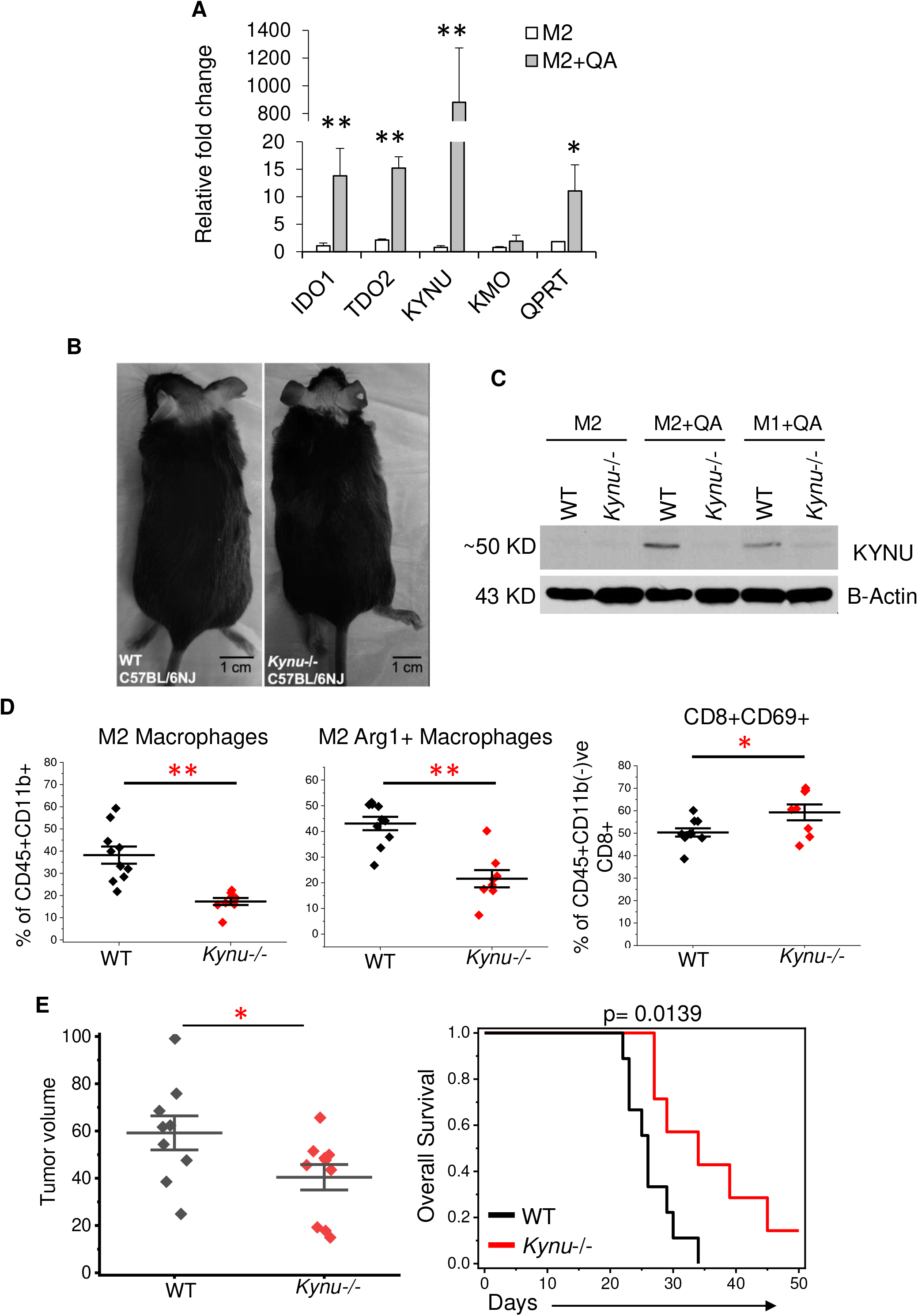
Targeting the downstream metabolism of tryptophan in GBM by inhibiting KYNU. (**A**) Macrophages obtained from C57BL/6 mice were polarized towards the M2 phenotype ± QA (20 μM), and RNA was isolated and evaluated for indicated genes using real-time PCR (n=3). (**B**) C57BL/6NJ *Kynu-/-* mice were generated, and phenotype compared to C57BL/6NJ wild type (WT) mice. (**C**) Macrophages obtained from C57BL/6NJ *Kynu-/-* and WT mice were polarized towards the M1 or M2 phenotype in ± QA, and indicated proteins were evaluated by western blot. (**D/E**) TRP tumors were grown orthotopically in C57BL/6NJ WT or *Kynu-/-* mice for (D) correlative and (E) survival studies. Specifically, a group of mice (n=10/group) were euthanized on d 21 and tumors harvested for immune profiling using flow cytometry. For survival studies, tumor volumes were measured using MR imaging on day 21 (n=10/group). *=p<0.05; **=p<0.005.

Next, using our *Kynu-/-* mouse model, we examined the influence of QA on the immune landscape of GBM *in vivo* and its potential to serve as a therapeutic target. As an initial investigation, cell lines derived from a GEMM^10, 32, 33^ were grown intracranially in WT and *Kynu-/-* mice, sacrificed on day 25, and tumors harvested for immunophenotyping and metabolite quantification. As expected, QA was nearly undetectable in tumors grown in *Kynu-/-* mice (**Fig. S9A).** No change in lymphocytes were observed when comparing WT and *Kynu-/-* mice; however, a significant decrease in total macrophages was observed in GBM tumors grown in *Kynu-/-* mice (**Fig. S9B**). Consistent with *in vitro* studies, a significant decrease in both M2 macrophages and M2 macrophages expressing the suppressive marker Arginase 1 (M2 Arg 1+) was demonstrated in GBM tumors grown in *Kynu-/-* mice (**Fig. 7D**), while no changes in MDSCs or Tregs were observed (**Fig. S9C**). Although no changes in numbers of CD4+ or CD8+ cells were observed, tumors grown in *Kynu-/-* mice had higher levels of CD8+CD69+ T cells (**Fig. 7D**) and CD8+ cells expressing Granzyme B (**Fig. S9D**), suggesting immune activation. Lastly, we performed imaging and survival studies to determine the potential of KYNU to serve as a therapeutic target. MR imaging performed on day 21 demonstrated a significant decrease in volume of tumors grown in *Kynu-/-* mice (**Fig. 7E**). GBM tumors grown in *Kynu-/-* mice had a significant improvement in median (25 d vs. 35 d) and overall survival, including ~50% of these mice still being alive when the last WT mice were euthanized, with several achieving long-term survival (**Fig. 7E**). Similar results were obtained when extending this line of investigation to the mouse GBM line GL261 (**Fig. S10**). Collectively, these studies support the potential of KYNU to serve as a novel therapeutic target to inhibit tumor growth by modulating the immune suppressive landscape of GBM.

## DISCUSSION

GBM continues to be an invariably fatal malignancy with limited treatment options. Despite continued advancements in surgical techniques, accuracy of radiation therapy delivery, and systemic therapy, median overall survival (OS) remains less than 2 years.^41^ Immune checkpoint agents represent a powerful new tool in the armamentarium of available therapies, revolutionizing how cancer is treated. Unfortunately, clinical gains demonstrated in a majority of other tumor types has not translated to improved outcomes in GBM patients.^42, 43^ One likely reason underlying this lack of clinical benefit of immune checkpoint inhibitors in GBM is the primary target of these agents may not play a dominant role in this tumor’s immune suppressive microenvironment. Rather than efforts designed to re-activate T cell antitumor activity by blocking the programmed cell death ligand-1 (PD-L1) axis or cytotoxic T-lymphocyte-associated protein 4 (CTLA-4), recent evidence suggests that TAMs play a more dominate role in the observed immunosuppression of GBM ^3, 44^. Macrophages can be polarized toward a pro-inflammatory/anti-tumor phenotype (M1), or an anti-inflammatory/immune suppressive phenotype (M2).^45, 46^ Tumors have been shown to have the capacity for polarizing macrophages toward the M2 phenotype, and the accumulation of these cells, in turn, helps tumors escape immune surveillance.^47^ M2 TAMs are found at higher percentages in GBM microenvironments compared to other tumor types, suggesting they play a critical role in the immune suppressive microenvironment found in these tumors ^44, 48^. Hence, there is considerable interest in developing therapeutic strategies designed to inhibit M2 TAMs or revert their polarization to enhance anti-tumor immunity in GBM. In these studies, we demonstrate the downstream metabolism of tryptophan, resulting in the accumulation of QA, represents a metabolic phenotype of GBM and uncover a novel role this metabolic pathway plays in sculpting the immune microenvironment by influencing macrophage polarization. As a major emphasis in the field has been the *upstream* metabolism of this pathway, namely, inhibiting the accumulation of the immune modulating metabolite Kyn by targeting IDO1 and/or TDO, these findings provide a previously undescribed window into the underlying biologic relevance of this pathway and its therapeutic implications. Specific to GBM, although increased expression of upstream enzymes IDO and TDO have been observed, along with the accumulation of Kyn,^12, 49^ targeting these pathways individually did not result in notable changes in the immune profile of these tumors or independent anti-tumor activity in preclinical models,^10, 13^ and unfortunately, as these investigations predicted, did not result in clinical benefit.^50, 51^ Based on our findings, targeting the *downstream* metabolism of tryptophan may have more relevance to the immune landscape of GBM and it is tempting to speculate, have a similar immune modulatory role in other types of cancers. Specifically, polarizing macrophages in the presence of QA resulted in nearly a 50% increase in the M2 phenotype, and importantly, conferred potent suppressive properties to these cells. Unexpectedly, QA also conferred an “M2-like” phenotype to macrophages polarized in conditions promoting M1 differentiation. In addition to expressing the M2 macrophage marker CD206, these cells functionally recapitulated M2 macrophages, both having suppressive properties and diminished phagocytosis capacity when tested using β-amyloid peptide. This offers the intriguing possibility that in addition to cancer, targeting this immune modulatory metabolite may have therapeutic implications in Alzheimer’s disease, which represents one of several neurodegenerative diseases where an accumulation of QA has been observed.^19^

Through global gene expression profiling, in addition to discovering a novel role the NMDA receptor plays in macrophage polarization, we uncovered an elegant mechanism linking QA-induced receptor activation to established pathways involved with M2 macrophage polarization. Specifically, through both chemical and molecular perturbation of this pathway coupled with ChIP analysis, we demonstrate QA leads to phosphorylation and degradation of the transcription factor Foxo1, which serves as an endogenous inhibitor of PPARγ expression through promoter binding. This resulted in both increased PPARγ expression, its transcriptional activation, and the observed increased M2 macrophage polarization. Further studies are warranted to more comprehensively establish the role of NMDA receptor activation and macrophage polarization and how this may apply to different pathophysiological states. We observed a potent positive feedback loop involving KYNU expression in macrophages cultured in the presence of QA, making it our lead candidate to inhibit the downstream metabolism of tryptophan. As agents are currently not commercially available to inhibit this enzyme, we generated a *Kynu-/-* knockout mouse model. These mice were phenotypically similar to wild-type mice and had identical immune profiles. Consistent with *in vitro* findings, tumors grown orthotopically in *Kynu-/-* mice demonstrated a ~50% reduction in M2 macrophages, a key cell thought to contribute towards the immune suppressive GBM microenvironment. Importantly, tumors implanted in these mice demonstrated decreased growth, resulting in improved survival, including several mice achieving long-term survival. As the link between tryptophan metabolism and cancer is well recognized,^13, 52^ and both preclinical and clinical studies designed to target the upstream metabolism of this pathway have thus far been disappointing,^51^ these findings offer an exciting opportunity to both revisit the biologic relevance of this pathway as it relates to oncogenesis and begin to identity compounds that can selectively target the downstream metabolism of tryptophan, thereby serving as a panel of novel immune modulatory agents that may be applied GBM, others cancers, and neurodegenerative diseases where this pathway has been implicated.

## METHODS

### Reagent and resource sharing

Further information and requests for resources and reagents should be directed to and will be fulfilled by the Lead Contact, Prakash Chinnaiyan.

### Human tumor samples and metabolomic/genomic profiling

Metabolomic profiling, gene expression profiling, GBM subtyping, O6-methylguanine-DNA methyltransferase (MGMT) promoter methylation, and isocitrate dehydrogenase 1 (IDH1) mutation analyses of patient-derived tumors were performed as previously described.^10, 28^

### Animal studies

All animal studies were carried out under protocols approved by the Institutional Animal Care and Use Committee at William Beaumont Research Institute (AAALAC-accredited facility) under animal protocols AL-15-09, AL-18-10, and AL-19-07). C57BL/6 mice were purchased from Charles River Laboratories (Wilmington, MA). C57BL/6NJ-*Kynu*-/+ and C57BL/6NJ were purchased from the Jackson Labs (Bar Harbor, ME) and bred at the Beaumont Research Institute animal facility to generate *C57BL/6NJ-Kynu-/-* mice. Mice with *Kynu* gene deletion were identified by Taqman probe PCR using primers (Integrated DNA Technologies, Coralville, IA) for *Kynu* deletion (F) 5’-CCTTGGCTATTTGTGATTGG-3’; *Kynu* wild type (F) 5’-GAAATTCCCTTGGCCTTCA-3’ and common (R) 5’-GAATTCTTCTGTCAGATGGAGTTACA-3’ along with *Kynu* deletion probe FAM-5’-AGCATAGTAACTGCGTGAGATCG-3’-Quencher and *Kynu* wild type probe TAMARA-5’-AGTGGGCCAAGATGTAAGTACC-3’-Quencher. Taqman PCR was performed using Vii7 real-time PCR using Jackson Laboratory’s recommended protocol for genotyping *Kynu* deletion. Animal experiments were conducted with an equal number of age and gender matched (6-10 weeks old male and female) mice.

### Cells culture and reagents

The TRP GBM line was generated from genetically engineered mice and cultured in MEM as previously described.^10, 32, 33^ Briefly, primary astrocyte lines were generated from a series of conditional GEM in which core GBM pathways were genetically targeted. Cre-mediated recombination *in vitro* lines express (1) a truncation mutant of SV40 large T antigen (T) from the human Gfap promoter that inactivates all 3 Rb family proteins, (2) a constitutively active KrasG12D mutant (R), and/or (3) a homozygous Pten deletion (P). GL-261 cells (murine GBM) were purchased from NIH’s DCTD tumor repository and cultured in DMEM. IC-21 cells (murine macrophage) were purchased from ATCC and cultured in RPMI-1640. Human iPSC-derived microglia cells were purchased from Fujifilm Cellular Dynamics, Inc. (Madison, WI) and cultured as described by Abud et. al.^53^ Briefly, cells were cultured in DMEM/F-12 media with N-2, B27, glutamax, ITS-G, and human insulin supplements (GIBCO/Thermo, Grand Island, NY) along with 1-thioglycerol, ascorbic acid and MEM non-essential amino acids (Millipore/Sigma, St. Louis, MO). In addition, cells were cultured in the presence of cytokines rhM-CSF (25 ng/ml), rhTGF-β1 (50 ng/ml), rhIL-34 (100 ng/ml), rhFractalkine (100 ng/ml) (PeproTech, Rocky Hill, NJ) and 100 ng/ml of human CD200 (Acro Biosystems, Newark, DE). On day five, rhIFN-γ (50ng/ml) and LPS (50ng/ml) were added to polarize these cells towards the M1 phenotype. Primary cell cultures of mouse splenocytes (T cells) was performed in RPMI-1640. All cell culture reagents were purchased from GIBCO/Thermo (Grand Island, NY) and Corning Life Sciences (New York, NY). Tryptophan, Kyn, QA, NAD+, and LPS (E. Coli O111:B4) were purchased from Sigma Aldrich (St. Louis, MO). All cytokines for murine macrophage polarization (mIL-4, mIL-13, mIFN-γ, hIFN-γ, hTGF-β, hIL-2, and mGM-CSF) were purchased from PeproTech (Rocky Hill, NJ). KMO inhibitor (UPF648), PPARγ inhibitor (GW9662), Foxo1 inhibitor (AS-1842856), PPARγ agonist (troglitazone), were purchased from Cayman Chemicals (Ann Arbor, MI). NMDAR agonist (NMDA) and antagonist (L-AP5) were purchased from Tocris Bioscience (Minneapolis, MN). GDC-0919 was obtained from Genentech (San Francisco, CA).

### Orthotopic tumor models

6-7-week old C57BL/6 mice were anesthetized and immobilized using a stereotactic frame, a midline scalp incision was made, and tumor cells (TRP: 1×10^5^ cells/mouse; GL-261: 1×10^4^ cells/mouse) suspended in 2 μl PBS were injected using a Hamilton syringe localized to specific coordinates (2 mm posterior to bregma, 2 mm mediolateral from the midline, and at a depth of 3 mm). Mice were imaged 6-8 days after implant using Multihance (Bracco Diagnostics, Cranbury, NJ, USA) diluted in sterile saline delivered via tail vein injection under anesthesia. MR imaging were performed as previously described.^54^ Briefly, images were acquired from anesthetized mice using a 3.0 T benchtop MRI (MR Solutions, Guildford, UK). Fast spin-echo T2 and T1 weighted images were acquired pre- and post-contrast injection. Tumor volumes were determined by delineating areas of contrast enhancement on the T1 weighted fast spin-echo sequence using the image processing PBAS tool in PMOD software (PMOD Technologies, Zurich, Switzerland). Mice were re-imaged after 2-3 weeks and followed until endpoint criteria were met. Similarly, orthotopic GBM tumors were implanted in C57BL/6NJ-*Kynu*-/- and C57BL/6NJ (WT) mice. For this experiment C57BL/6 mice were injected orthotopically with TRP cells (1×10^5^ /mouse). Mice were imaged for tumor implants on day 8. A group of mice were treated with the IDO inhibitor (GDC-0919) using oral gavage twice a day (200 mg/kg) for two weeks (6 days a week) starting on day 8 following tumor implant. Tumors were isolated on day 25 to analyze QA and KYN levels.

### Immune cell polarization

#### Macrophage polarization

Bone marrow cells were isolated from the indicated mouse line and cultured in GM-CSF (40ng/ml) for 5-7 days to generate M0 cells. Cells were then cultured with IL-4 (20ng/ml) and IL-13 (20ng/ml) for M2 polarization or IFN-γ (50ng/ml) and LPS (100ng/ml) for M1 polarization (2 d). Bone marrow cells were cultured for 5 days in GM-CSF (40ng/ml) and IL-6 (40 ng/ml) to generate MDSCs. *Microglial cell polarization:* Microglial cells were isolated from brains of C57BL/6 mice using the Percoll density gradient centrifugation method ^55^. Microglial cells were cultured in GM-CSF (40ng/ml) for 5 days to generate M0-MDMs (microglial-derived macrophages). Cells were then cultured with IL-4 (20ng/ml) and IL-13 (20ng/ml) for MDM2 polarization or IFN-γ (50ng/ml) and LPS (50ng/ml) for MDM1 polarization (2 d). *iTreg polarization:* Magnetic bead sorted naïve CD4+ T cells were isolated from C57BL/6 mice splenocytes. CD4^+^ T cells were polarized to CD4^+^ FoxP3^+^ using rhTGF-β (30 ng/ml) and rhIL2 (300 IU/ml) and activated using plate-bound anti-CD3 (1μg/ml; clone-145-2C11) and anti-CD28 (5μg/ml; clone-37.51) antibodies (Biolegend; San Diego, CA) for 4 days.

### Immunohistochemical staining

TMA sections were obtained from US Biomax (Derwood, MD) and comprised of human astrocytoma grades II, III along with 30 GBM tumors and 9 normal brain sections. Immunohistochemical staining was performed on FFPE sections deparaffinized and rehydrated using a Leica Autostainer, followed by Tris base (pH 9.0) dependent protein antigen retrieval (Vector Labs, Burlingame, CA) at 95°C. Slides were washed with PBS, incubated in H2O2 (3% in PBS) for 10 min, washed, and blocked with 5% goat serum in PBS-T (PBS with 0.3% Triton X100) for 30 minutes. QA antibody (clone 4E11-G3, 1:500 ImmuSmol, France) was diluted 1:500 in antibody diluent solution (Thermo, Grand Island, NY). Sections were incubated with QA antibody at 4°C overnight, washed with PBS, and incubated with an HRP-conjugated secondary antibody at room temperature for 1 hour. Chromogen DAB (Vector Labs, Burlingame, CA) was applied for 8-10 min after washes. Slides were counter stained using a Leica Autostainer. A section was included in which only the HRP-conjugated secondary antibody was used to serve as a negative control. Sections were imaged using a Aperio Imagescope (Leica Biosystems, Buffalo Grove, IL) at magnification fields (x400). QA staining was quantified by dividing staining intensity into 4 subgroups: strong (3), intermediate (2), low (1), or absent (0). Intensity was multiplied by a factor of 2 if > 10 positive foci were observed in each field.^56^ Three fields were observed for each TMA sample.

### T cell culture

Primary splenocytes were obtained from C57BL/6NJ or C57BL/6NJ-*Kynu*-/- mice. Cells were analyzed for CD4+ T cells, CD8+ T cells, NK1.1 (NK cells), B cells (CD19) and Tregs (CD4+CD25+FoxP3+). Magnetic-bead sorted naïve T cells (CD4+ and CD8+ T cells) were activated with plate-bound anti-CD3 (1μg/ml; clone-145-2C11) and anti-CD28 (5μg/ml; clone-37.51) antibodies (Biolegend) for 3 days. CD4+ and CD8+ T cells were analyzed for markers of activation (CD69+) and proliferation (Ki67). Activated CD4+ and CD8+ T cells were stimulated with a cell activation cocktail (Biolegend; CA PMA/Ionomycin with Brefeldin A) overnight and stained for IFN-γ and TNF-α after fixation and permeabilization using the eBioscience/Thermo kit (Grand Island, NY). Cells were and acquired using the FACS Canto II flow cytometer.

### CD8 T cell proliferation/suppression

CD8^+^ T lymphocytes were sorted using Dynabead FlowComp Mouse CD8 Kit (Grand Island, NY). Cells were stained with CFSE CellTrace (GIBCO/Thermo; Grand Island, NY) and activated with plate-bound anti-CD3 (1μg/ml; clone-145-2C11) and anti-CD28 (5μg/ml; clone-37.51) antibodies (Biolegend). For suppressing proliferation, polarized M2 macrophages were added at varying concentrations with and without Kyn or QA. CFSE dilution on CD8^+^ T cells was analyzed 3 days after activation using a FACS Canto II flow cytometer (Becton Dickinson, Mountain View, CA). CFSE dilution was analyzed on CD8^+^ T cells after 3 days of activation. CFSE stained cells were also stained for Granzyme B, (clone 29F.1A12).

### Flow cytometry

Tumors were collected from mice and were dissociated using DNase I and Collagenase IV (Sigma Aldrich, St. Louis, MO). Cells were stained for M2 macrophages (CD45+F4/80+CD11b+CD206+) (CD45+F4/80+CD11b+CD206+IL-4Ra), M2 cells with Arginase 1 (CD45+F4/80+CD11b+CD206+Arg1+), M1 macrophages (CD45+F4/80+CD11b+CD80hi), MDSCs (CD45+Gr1+CD11b+), Tregs (CD4+CD25+FoxP3+), CD8 T lymphocytes (CD45+CD11b-CD8+), activated CD8 cytotoxic T lymphocytes (CD45+CD11b-CD8+CD69+), granzyme B+ CD8 T lymphocytes (CD45+CD11b-CD8+GzmB+), CD4 T lymphocytes (CD45+CD11b-CD4+). Antibodies were purchased from Biolegend or eBioscience/Thermo. Clones for antibodies used for flow cytometry are as indicated: CD45 (30-F11), F4/80 (BM8), CD11b (M1/70), CD206 (C068C2), Arg 1 (A1exF5), IL4Ra (JAMA-147), CD80 (16-10A1), Gr1 (RB6-8C5), FoxP3 (150D/E4), CD8 (53-6.7), CD4 (Gk1.5), CD25 (PC61), CD69 (H1.2F3), GzmB (QA16A02). Fluorochrome-linked anti-NMDAR1 antibody (N308/48) was obtained from Abcam (Waltham, MA). Fixation and permeabilization was performed using the eBioscience/Thermo kit. For analysis of the phagocytotic ability of cells, green (HiLyte Fluor 488) florescent β-amyloid (1 - 42) peptide (Anaspec, Fremont, CA) was used. Macrophages/microglial cells were pulsed with 400 nM of florescent peptide for 16 hours, followed by washing and removal of the soluble florescent probe. Cells were then stained for surface markers (CD45, F4/80, CD11b, CD206, or CD80). All samples were analyzed on a FACS Canto II flow cytometer (Becton Dickinson; Mountain View, CA). Analysis of flow cytometry data was performed using FlowJo V10 software (FlowJo, LLC; Ashland, OR).

### Aryl hydrocarbon receptor (AhR) activity reporter assay

AhR activation was analyzed using AhR reporter cells (INDIGO Biosciences, PA) according to the manufacturer’s protocol.

### Chromatin immunoprecipitation (ChIP)

ChIP was performed using a ab500-ChIP kit (Abcam, Cambridge, MA) according to the manufacturer’s protocol. Briefly, cells were cross-linked in 1.1% formaldehyde and then resuspended in cell lysis buffer followed by sonication for DNA fragmentation (200-1000bp). Chromatin was sonicated and then incubated with a polyclonal antibody against PPARγ (81B8 and C26H12) or FoxO1 (C29H4) Cell Signaling Technology (Danvers, MA) for immunoprecipitation. Antibodies for histone H3 was used as a positive control and IgG as a negative control. ChIP-enriched DNA was further purified and quantified by real-time PCR for the PPARγ gene and normalized to input DNA control samples using PPARγ primers F-5’-CCACTGGTGTGTATTTTACTGC-3’ and R5’-AAAATGGTGTGTCATAATGCTG-3’ ^57^. RLP30 primers were obtained from Cell Signaling Technology and used as a control.

### Western Blot

Western blot was performed using methods previously described.^10^ PPARγ (C26H12), Foxo1 (C29H4), pFoxo1 (Ser256) (#9461), and tubulin (#2144) antibodies were obtained from Cell Signaling Technology. Mouse KYNU (polyclonal), tubulin, and actin antibodies were obtained from Thermo Fisher.

### Real-time qPCR

Total RNA was extracted from pellets of polarized macrophages, and mRNA was extracted using RNA mini kit (BioRad, Hercules, CA), and cDNA was generated from total RNA using iScript cDNA Synthesis Kit (BioRad, Hercules, CA). SYBR Green (Biorad, Hercules, CA) was used for quantitative PCR using Vii7 real-time PCR (Grand Island, NY). Primers were synthesized using a predesigned qPCR primer from IDT (Coralville, IA). Expression mRNA was normalized to Actb and HPRT.

### Murine microarray and data analysis

Murine M2 polarized cells were gated for CD45+CD11b+F4/80+CD206+ and flow-sorted using FACS Aria II (Becton Dickinson; Mountain View, CA). Cells were more than 98% pure. Total RNA was isolated from these cells using the Qiagen all prep kit (Hilden, Germany; cat # 80204) and quantified using a Nanodrop. Integrity of RNA preparation was confirmed with the Agilent RNA 6000 Nano Kit (cat # 5067-1511). 500 ng of total RNA was used for Clariom D mouse Transcriptome Array 1.0 (MTA 1.0.). Raw data were processed using the Affymetrix expression console, and further data analyses were performed using the Affymetrix transcriptome analysis console. Raw data were normalized using gene-level RMA Sketch, and bi-weight average signal (log2) values were used for analysis. To determine the association between various mRNA gene expression, we performed hierarchical clustering with log2 transformed normalized data using Euclidean distance and Ward scaling using MetaboAnalyst 5.0 ^58^. Fold change was calculated and used in generating the volcano plot (log2 fold change on the x-axis and negative log10 of p-value on the y-axis). Using the top 24 genes from the volcano plot, we performed protein-protein interaction analysis using Pathway Commons version 12 ^59^. Transcriptions factors (AP1, AP4, FOXO1, FOXO4, PPARγ, OCT1, AKT1, NFKB, STAT1, IRF1, IRF2, IRF7) associated with target genes were integrated into the analysis to identify interactive pathway.

### PPARγ transcriptional activation

PPARγ Transcription Factor ELISA Assay was used to analyze PPARγ binding activity according to the manufacturer’s protocol (Caymen Chemicals, Ann Arbor, MI).

### PPARγ silencing

PPARγ silencer select (assay ID-s72013) and non-targeting (scrambled si; Thermo Fisher) was used for transient knockdown. IC-21 cells were cultured in ±QA for 6 days in RPMI-1640 along with GM-CSF (40ng/ml). Cells were then transfected with siRNAs using Lipofectamine 2000 (Thermo Fisher Life Technologies) for 8 h in Opti-MEM media (Thermo Fisher). Cells were re-suspended in macrophage media containing IL4 and IL13 (20 ng/ml) for 24 h to polarize towards the M2 phenotype. Cells were then transfected with siRNAs using Lipofectamine 2000 (Thermo Fisher Life Technologies) for 8 h in Opti-MEM media (Thermo Fisher) and re-suspended in macrophage conditioning media.

### Quinolinic acid quantification

QA was quantified using ELISA (Abclonal, Woburn, MA). Tryptophan (Trp), QA, and KYN were analyzed using or gas-chromatography–mass spectrometry ^60, 61^. Briefly, weighted tumor tissues or an equal number of pelleted cells were treated with equal parts of ice-cold methanol and water. Subsequently, the cell pellet was homogenized. Ice-cold chloroform was added to the cell pellets, followed by centrifugation to get a phase separation. The aqueous phase was air-dried and derivatized (75°C) using PFP (2,2,3,3,3-pentafluoro-1-propanol) and PFPA (pentafluoropropionic anhydride), and the derivatized samples were subsequently dried using a stream of nitrogen. Dried samples were dissolved in ethyl acetate and acquired using Agilent 7000C triple quad mass spectrometer and Agilent 7890 GC (Agilent, Santa Clara, CA). 1μL sample was injected using Agilent 7693 auto-sampler injector operated in electron capture negative ionization (ECNI) mode. Trp, QA and KYN were quantified by selected ion monitoring of QA (431 m/z), KYN (454 m/z), and Trp (608 m/z). Data were analyzed using MassHunter version B (Agilent, Santa Clara, CA). Similar cell pallets or tissues were spiked with different concentrations of QA and compared with test samples to get an approximate quantification of QA.

### Statistical analysis

Comparisons across two different groups were performed on original data using 2 tailed student’s T test. A log-rank test was used for survival analyses. All statistical analyses were performed using Origin Pro 2019 software (Origin Lab Corporation; Northampton, MA).

## ACKNOWLEDGMENTS

This work was supported in parts by the National Institute of Health (NIH)/National Institute of Neurological Disorders and Stroke (NINDS) grants R01NS110838, R21NS090087, American Cancer Society grant RSG-11–029-01, and Bankhead-Coley Cancer Research Program and Cancer Research Seed Grant Awards from Beaumont Health to Prakash Chinnaiyan. CRM is supported by NIH/NCI (R01CA204136). The authors sincerely acknowledge help provided by Barbara Pruetz (Beaumont Biobank) in the standardization of IHC for QA.

## AUTHOR CONTRIBUTIONS

Study’s conception and design-PK, SK, and PC. Experimentation and data acquisition - PK, SK, YZ, AP, and KLB. Analysis and data interpretation - PK, SK, YZ, AP, KLB, and PC. Writing of the article -PK, SK, PC, and CRM. All authors contributed to review and editing the article.

## COMPETING INTERESTS

All authors have reviewed and approved the manuscript. Authors declare no competing interests.

## Notes

### Competing Interest Statement

The authors have declared no competing interest.

