## Supplemental Figures for "Quinolinate Promotes Immune Tolerance through Macrophage Polarization in Glioblastoma"

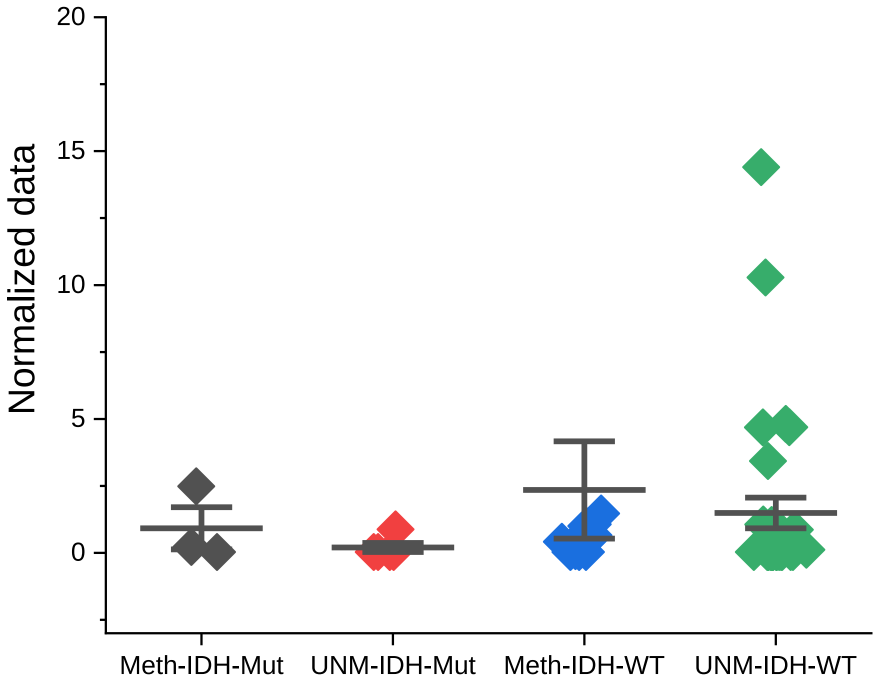


**Supplementary Figure 1**

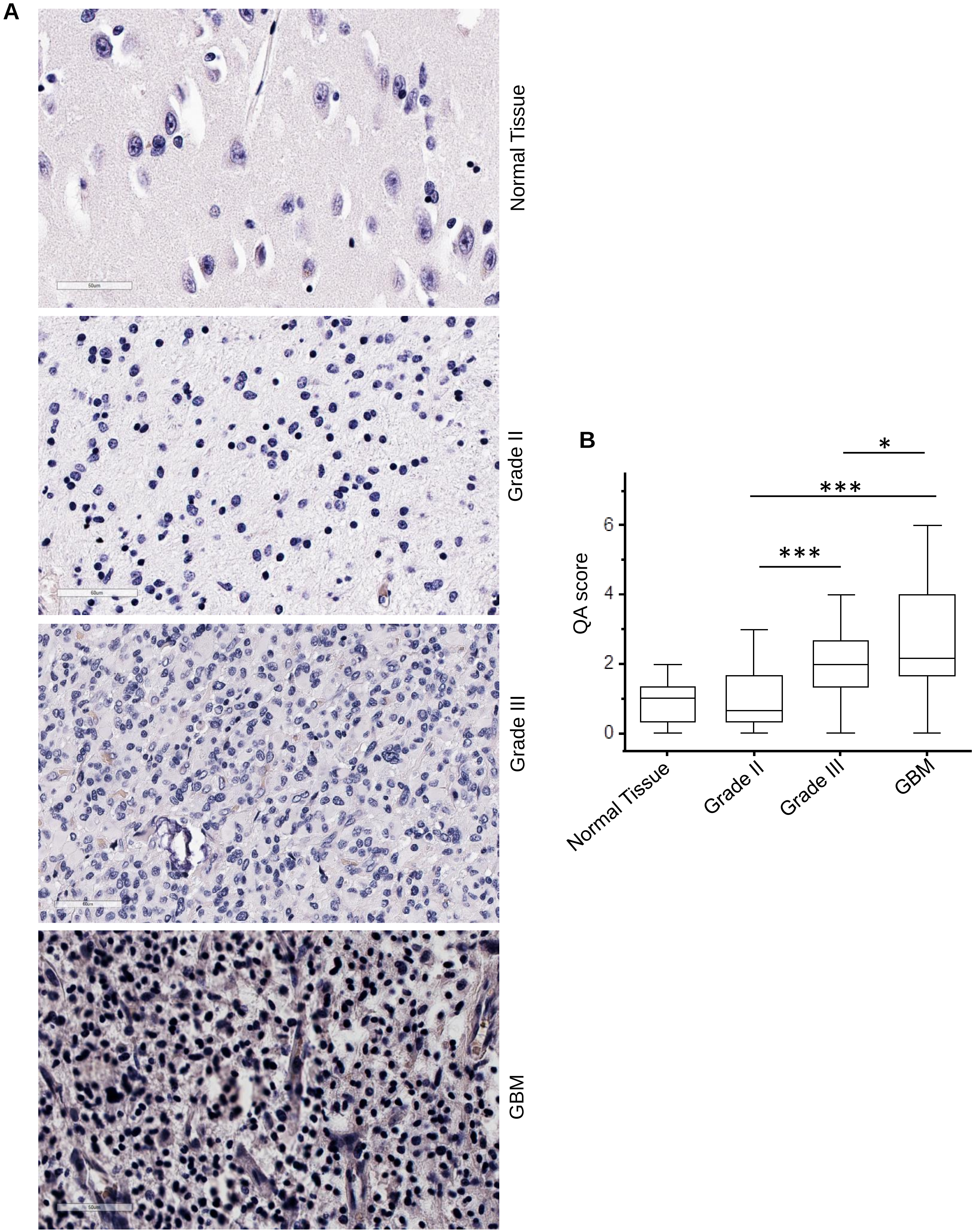


**Supplementary Figure 2**

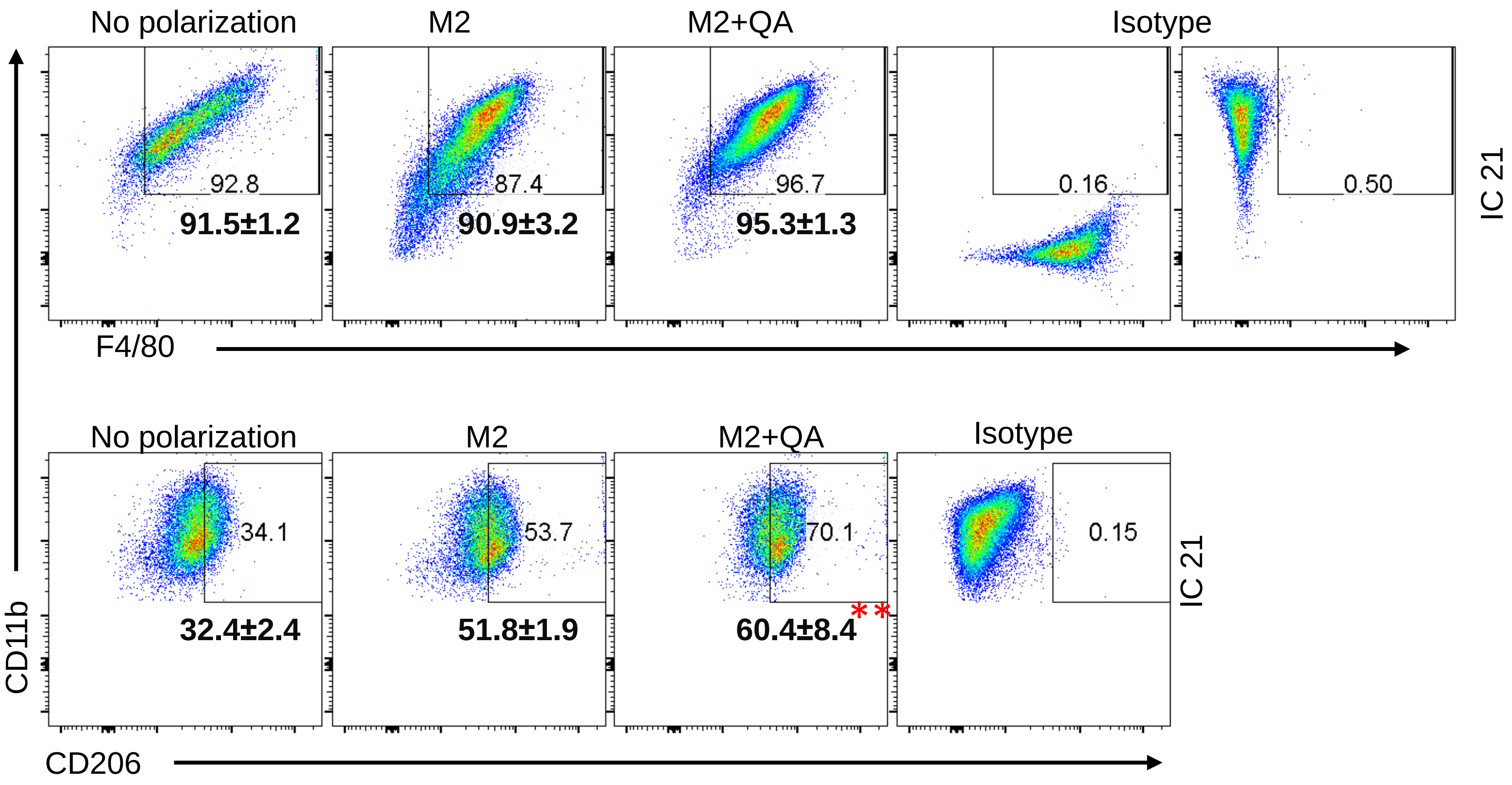


**Supplementary Figure 3**

**A**

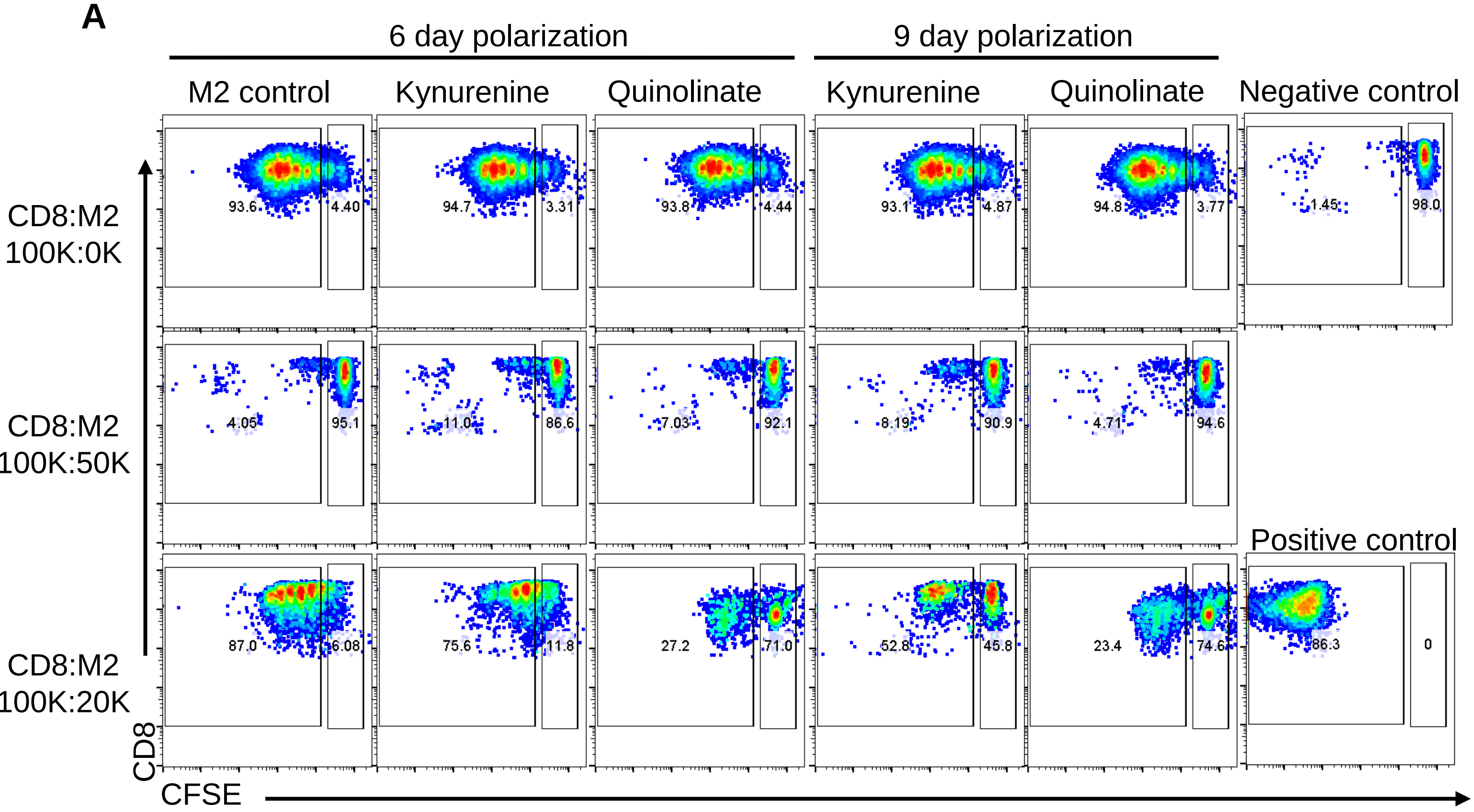


**B**

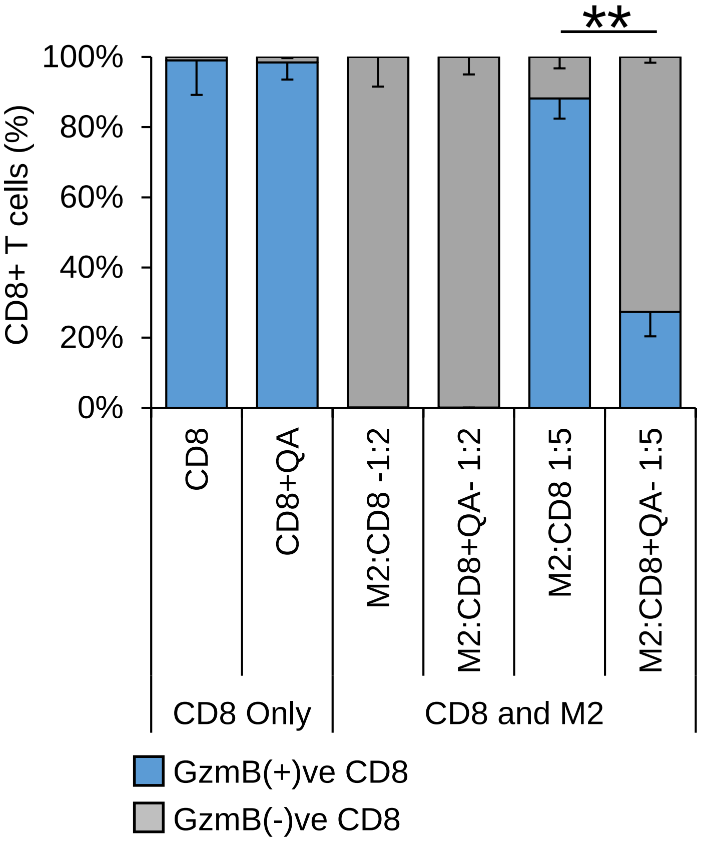


**Supplementary Figure 4**

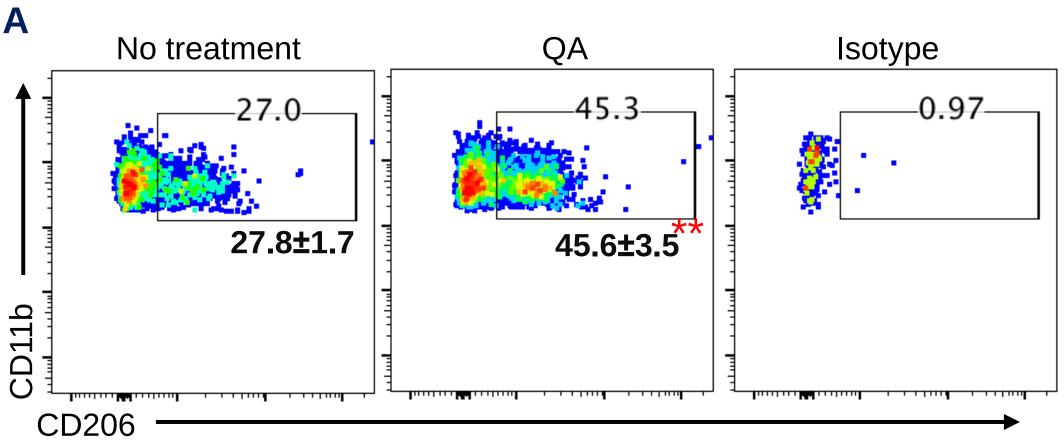


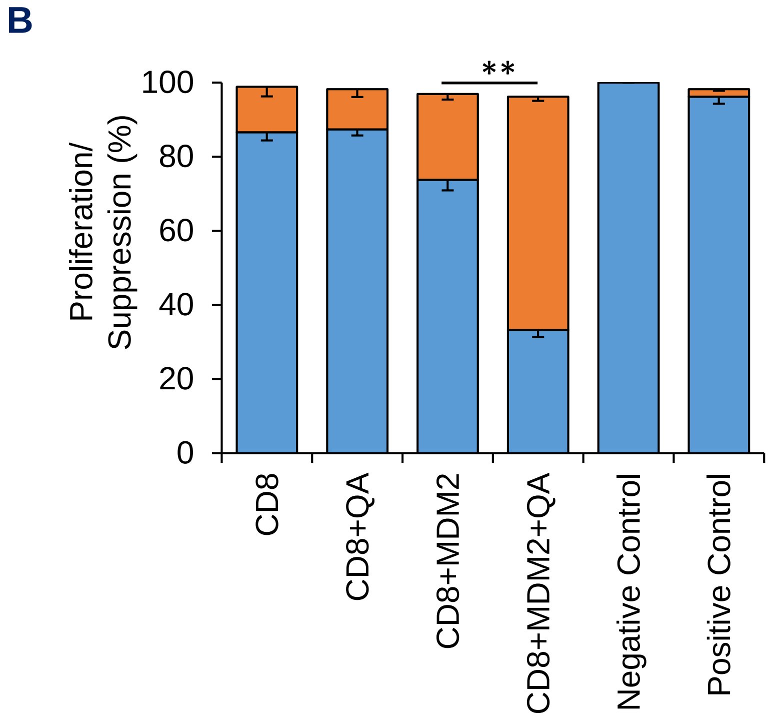


**Supplementary Figure 5**

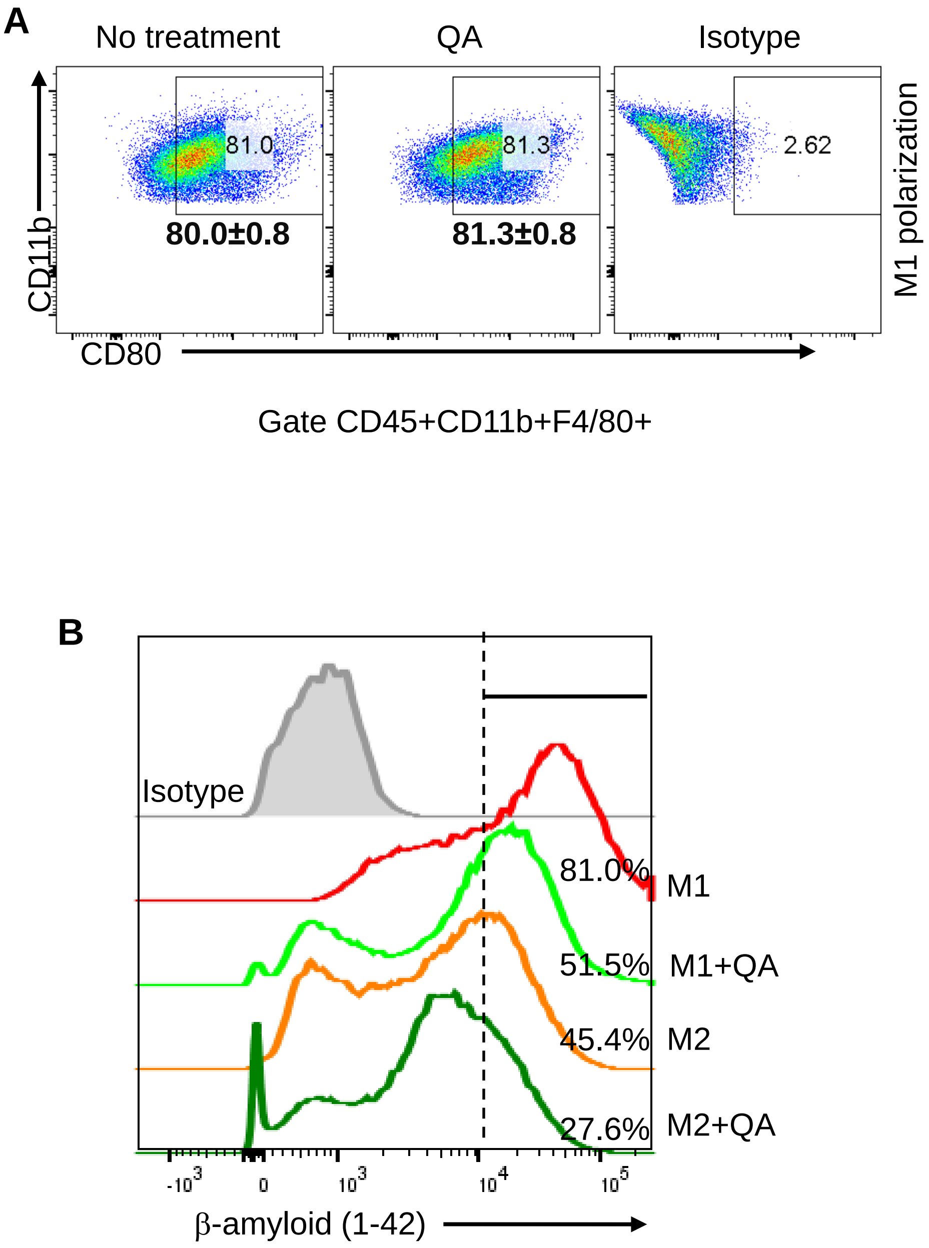


**Supplementary Figure 6**

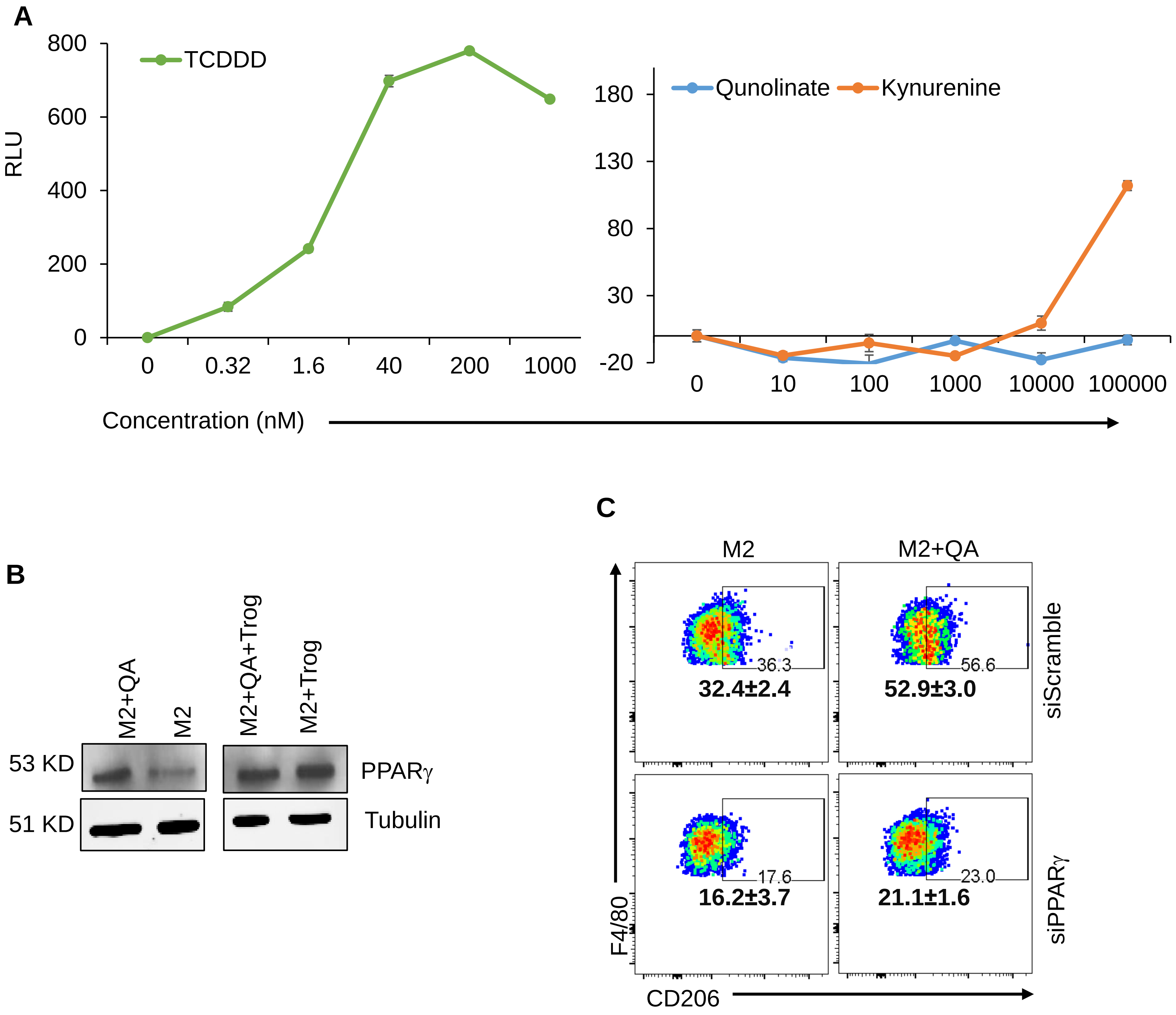


Supplementary Figure 7

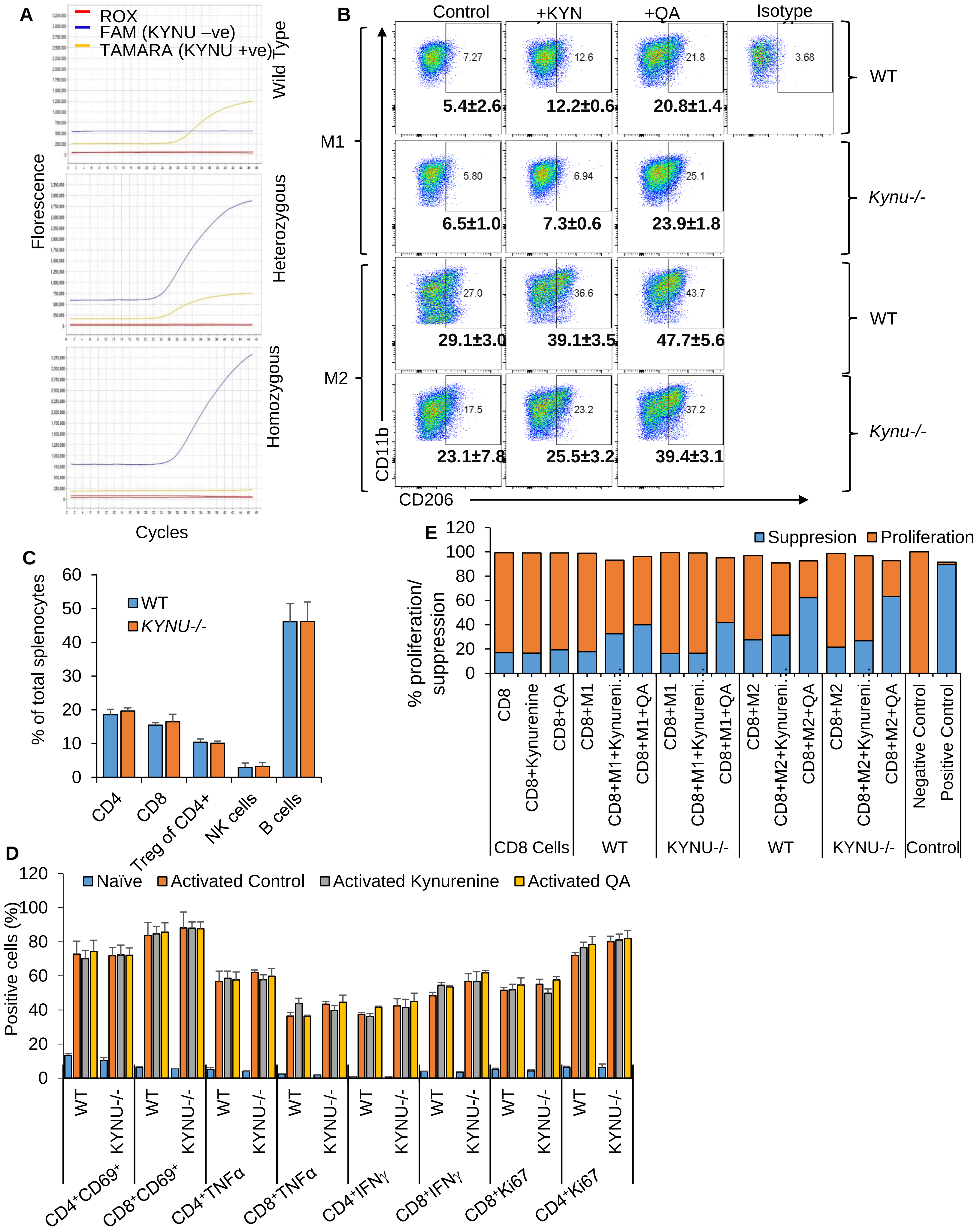


**Supplementary Figure 8**

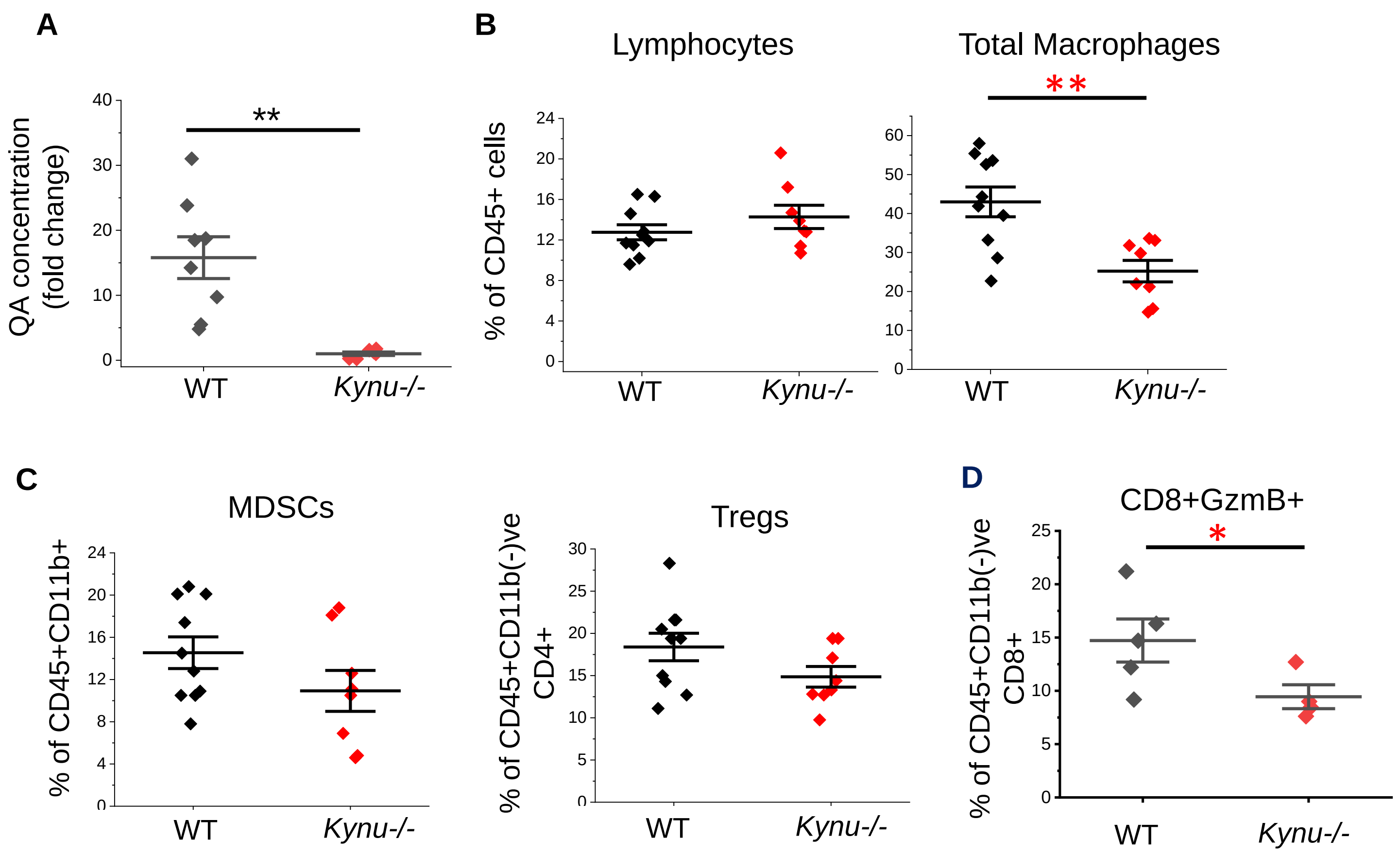


**Supplementary Figure 9**

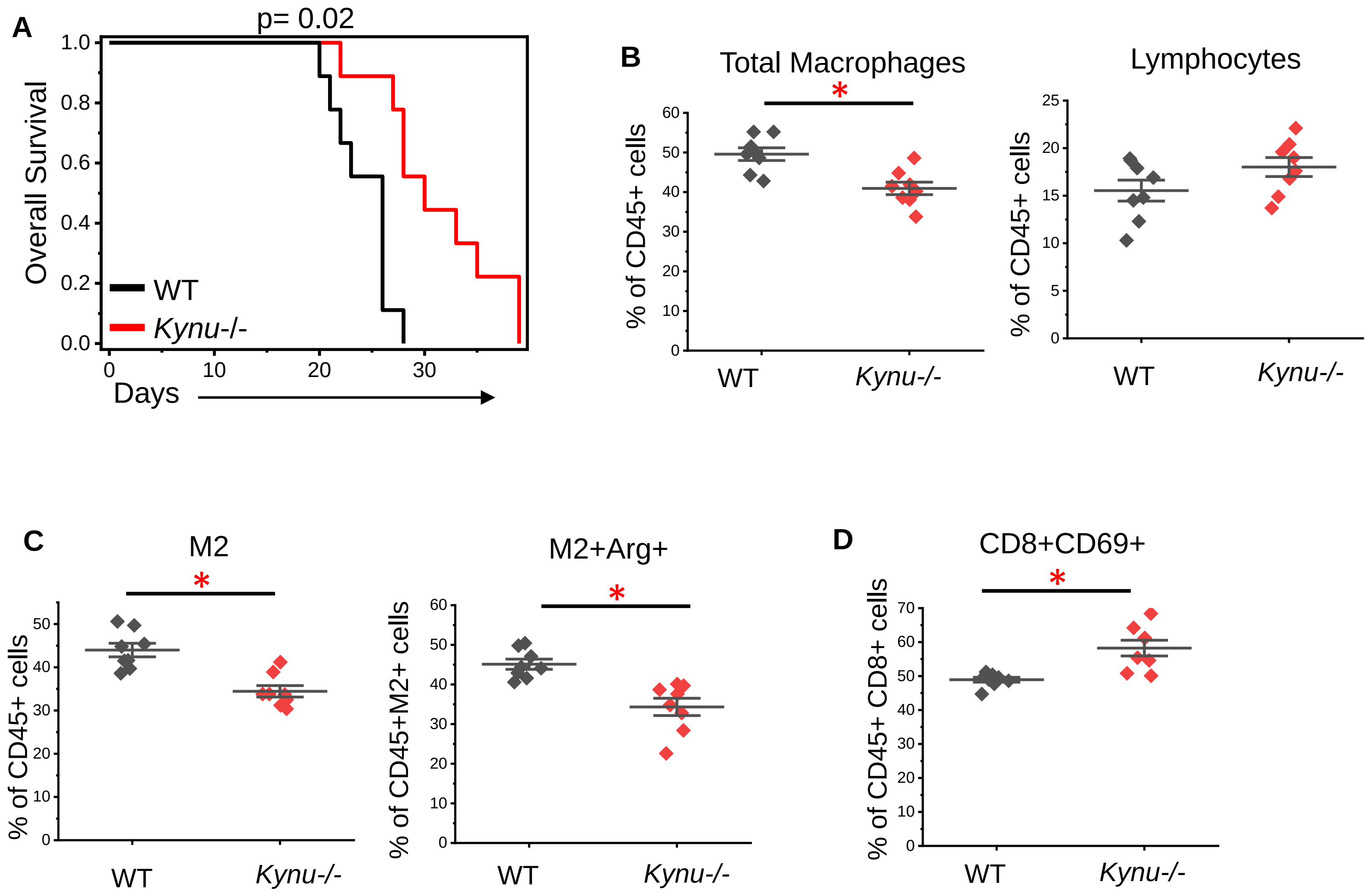


**Supplementary Figure 10**
