## Supplemental Figure Legends for "Quinolinate Promotes Immune Tolerance through Macrophage Polarization in Glioblastoma"

### SUPPLEMENTARY FIGURE LEGENDS

**Supplementary Figure 1:** The relative accumulation QA as a function of IDH mutation (wild-type [WT] vs. mutant [mut]) and MGMT methylation (methylated [meth] vs. unmethylated [UNM]) status in GBM (n=56).

**Supplementary Figure 2:** Immunohistochemical staining of QA in human astrocytoma. (A) Representative depiction of QA immunohistochemical staining and (B) bar graph depicting staining score in normal human brain tissue (n=7), grade II astrocytoma (n=16), grade III astrocytoma (n=12), and GBM (n=31) human tumors. Each tumor was analyzed in triplicate. *=p<0.05; **=p<0.005.

**Supplementary Figure 3:** The murine macrophage cell line IC-21 was cultured in GM-CSF (40ng/ml) for 6 days ± QA (20 μM). On day 6, cells were re-suspended in IL4 and IL13 (20 ng/ml) for 24 h to polarize towards the M2 phenotype. Cells were analyzed for M2 macrophage markers (CD45+CD11b+F4/80+CD206+). Numbers represent mean±SD from 3 different experiments. **=p<0.005.

**Supplementary Figure 4:** This figure is complementary to Fig. 2D, evaluating the suppressive ability of M2 macrophages when evaluated using varying ratios of CD8 T cells. M2 cells were polarized ± Kyn or QA for 6 or 9 d. Splenocytes from C57BL/6 mice were used for isolating $CD8^+$ T cells using magnetic bead sorting. CFSE labeled $CD8^+$ T cells were activated using plate-bound anti-CD3/CD28 antibody for three days in the presence or absence of M2 cells ± Kyn or QA. **(A)**

The proliferation of $CD8^+$ T cells is demonstrated by CFSE dilution. CFSE labeled $CD8^+$ T cells without stimulation (anti-CD3/CD28 antibody) were used as a positive control of suppression. Unlabeled $CD8^+$ T cells were used as a negative control for proliferation. Data is representative of 2 independent experiments. **(B)** $CD8^+$ T cells were analyzed for granzyme B (GzmB). The bar graph represents mean±SD from 2 different mice performed in duplicate.**=p<0.005.

**Supplementary Figure 5: (A)** Murine microglia derived M2-like macrophages (MDM2) generated from the microglia isolated from brain of C57BL/6 mice were cultured in GM-CSF (40ng/ml) for 7 days in ± QA and polarized to the M2 phenotype with IL4 and IL13 (20 ng/ml) for 48 h to polarize towards the M2-like phenotype. Cells were analyzed for M2 macrophage-specific markers (CD45+F4/80+CD11b+CD206+). Numbers represent mean±SD from 2 experiments performed in duplicate. **=p<0.005. **(B)** $CD8^+$ T cells (bead sorted from splenocytes of C57BL/6 mice). CFSE labeled $CD8^+$ T cells were activated using plate-bound anti-CD3/CD28 antibody for three days in the presence or absence of MDM2 cells ±QA. **=p<0.005.

**Supplementary Figure 6: (A)** M0 macrophages generated from the bone marrow of C57BL/6 mice were cultured in GM-CSF (40ng/ml) for 6 days ± QA (20 μM) and polarized to the M1 phenotype with LPS (100 ng/ml) and IFN- $\gamma$ (50ng/ml) for 24-36 h ± QA. Cells were analyzed for M1 macrophage-specific markers (CD45+F4/80+CD11b+CD80hi). **(B)** Murine M0 macrophages were cultured in the presence of GM-CSF and polarized towards the M1 (LPS+IFN$\gamma$) or M2 (IL4+IL13) phenotype ± QA. Cells were pulsed with green florescent $\beta$-amyloid (1-42) peptide and analyzed for phagocytosis of this peptide at 16 h. Macrophages were analyzed for the presence of green florescent $\beta$-amyloid (1-42) peptide by flow cytometry after gating on M1 or M2 specific markers. Results are representative of 2 independent experiments.

**Supplementary Figure 7: (A)** AhR reporter cells (firefly luciferase, 62 kD protein) were treated with varying concentrations of Kyn, or QA for 24 hours. 2,3,7,8-tetrachlorodibenzo-p-dioxin (TCDD - ligand of AhR activation) was used a positive control. Luciferase was measured using a luminescence reader as relative light units (RLUs). **(B)** Macrophages obtained from C57BL/6 mice were polarized towards the M2 phenotype ± QA (20 μM) or the PPARγ agonist troglitazone (Trog; 5 μM) and evaluated for the indicated proteins by western blot. **(C)** The murine macrophage cell line IC-21 was cultured in GM-CSF (40ng/ml) for 5 days ± QA. On day 5 siRNA was used to perform knockdown of PPARγ (siPPARγ). On day 6, cells were re-suspended in IL4 and IL13 (20 ng/ml) for 24 h to polarize towards the M2 phenotype. Cells were analyzed for M2 macrophage markers (CD45+CD11b+F4/80+CD206+). Numbers represent mean±SD from 3 different experiments.

**Supplementary Figure 8: (A)** C57BL/6-NJ and *KYNU-/-* mice genotyped using the Taqman PCR assay. FAM probe was used to detect knockout of KYNU, TAMARA Taqman probe detected the wild type KYNU strand. ROX was used as a background control. **(B)** C57BL/6-NJ and *Kynu-/-* mice were isolated and polarized to M1 or M2 phenotype in ± QA or ± Kyn for 9 days. Cells were analyzed using flow cytometry for the M2 macrophage marker (CD45+CD11b+F4/80+CD206+). Numbers represent mean±SD from 3 different experiments. **(C)** Naïve splenocytes were isolated from C57BL/6-NJ (WT mice) and *Kynu-/-* mice and analyzed for $CD4^+$ and $CD8^+$ T cells, T regulatory cells ($CD4^+FoxP3^+CD25^+$), NK cells (NK1.1+), and B cells (CD19+). The bar graph represents mean±SD from 3 different experiments. **(D)** CD4+ T cells and CD8+ T cells were magnetic bead sorted from C57BL/6-NJ (WT mice) and *Kynu-/-* mice splenocytes and were activated using a plate-bound anti-CD3/CD28 antibody for three days. $CD4^+$ and $CD8^+$ T cells were analyzed for activation marker (CD69) and proliferation marker (Ki67). Activated $CD4^+$ and

$CD8^+$ T cells were incubated with PMA/Ionomycin+Protein Transport Inhibitor (Brefeldin A) for 16 hours and analyzed for TNFα and IFNγ. **(E)** Macrophages obtained from C57BL/6-NJ and *Kynu-/-* mice were polarized ± Kyn or ±QA (20 μM) for 9 days. Splenocytes from C57BL/6-NJ mice were used for isolating $CD8^+$ T cells using magnetic bead sorting. CFSE labeled $CD8^+$ T cells were activated using plate-bound anti-CD3/CD28 antibody for three days in the presence or absence of macrophages ± Kyn or QA. The proliferation of $CD8^+$ T cells is demonstrated by CFSE dilution. CFSE labeled $CD8^+$ T cells without stimulation (anti-CD3/CD28 antibody) were used as a positive control of suppression. Unlabeled $CD8^+$ T cells were used as a negative control for proliferation.

**Supplementary Figure 9:** TRP tumors were grown orthotopically in C57BL/6-NJ (WT) mice or *Kynu-/-* mice. Mice (6-9/group) were euthanized on day 21 of the tumor implant. Tumors were harvested and used for **(A)** QA analysis using ELISA and **(B/C)** immune phenotyping. **(B)** Orthotopic TRP tumors were harvested from C57BL/6-NJ (WT) mice or *Kynu-/-* mice were used for comprehensive flow cytometric analysis of lymphocytes and total macrophages; **(C)** along with MDSCs (CD45+CD11b+Gr1+); Tregs (CD45+CD4+FoxP3+CD25+) and **(D)** Granzyme B + CD8+ T cells. *=p<0.05; **=p<0.005.

**Supplementary Figure 10: (A)** GL261 murine GBM tumors were grown orthotopically in C57BL/6NJ WT or *Kynu*-/- mice for survival studies (n=6/group). **(B-D)** In a similar experiment, mice were euthanized on d 21 and tumors harvested for immune profiling using flow cytometry (n=8/group), including total macrophages (CD45+CD11b+F4/80+), lymphocytes (CD45+CD11b-ve); M2 macrophages (CD45+CD11b+F4/801+CD206+); Arginase+ M2 macrophages (CD45+CD11b+F4/801+CD206+Arg1+), and activated CD8+ T cells

(CD45+CD8+CD69+). Line between the data points represents mean and whisker represents SE. *=$p<0.05$
